# Adaptation of α-synuclein fibrils following multiple system atrophy transmission to mice

**DOI:** 10.64898/2026.05.06.723086

**Authors:** Mylan Mayer, Chase R. Khedmatgozar, Gianna Zinnen, Matthew P. Frost, Patricia M. Reis, Sara A. M. Holec, Melissa Dexter, Arthur A. Melo, Eric Tse, Gregory E. Merz, Amanda L. Woerman

**Affiliations:** Institute for Neurodegenerative Diseases, Weill Institute for Neurosciences, University of California San Francisco, San Francisco, CA, USA; Department of Microbiology, Immunology, & Pathology and Prion Research Center, Colorado State University, Fort Collins, CO, USA; Department of Biology, University of Massachusetts Amherst, Amherst, MA, USA; Cell and Molecular Biology Graduate Program, Colorado State University, Fort Collins, CO, USA; Department of Neurology, University of California San Francisco, San Francisco, CA, USA; Department of Pharmaceutical Chemistry, University of California San Francisco, San Francisco, CA, USA

## Abstract

Synucleinopathies are a group of neurodegenerative diseases characterized by the presence of misfolded α-synuclein inclusions which cause progressive disease by spreading throughout the brain in a prion-like manner. Throughout the neurodegenerative disease field, the ability of a single protein to give rise to multiple distinct clinical disorders is explained by the strain hypothesis, or the idea that the misfolded protein conformation determines the resulting disease. This was initially shown using transmission studies in cell lines and mouse models; more recently cryo-electron microscopy (cryo-EM) validated this idea by identifying distinct α-synuclein filament folds in brain tissues from patients with Parkinson’s disease, multiple system atrophy (MSA), and juvenile-onset synucleinopathy. However, very little is known about the α-synuclein filament structures that form in animal models of these disorders, and thus their relevance to human disease and suitability as models for therapeutic development remains a question. Here we report the first atomic resolution cryo-EM structures of α-synuclein fibrils from an MSA patient sample before and after transmission to a transgenic mouse model of disease. Our findings indicate that while distinct adaptations occur during fibril replication in the mouse host, key structural facets are maintained, validating the merits of this transmission model for supporting preclinical research on MSA.

## INTRODUCTION

Synucleinopathies are a group of neurodegenerative diseases, including Parkinson’s disease (PD) and multiple system atrophy (MSA), that are neuropathologically defined by the presence of misfolded inclusions of the protein α-synuclein in the brain. Over a decade of research investigating α-synuclein biology in transmission models (e.g., cell lines and transgenic mice) has repeatedly demonstrated that the protein behaves in a prion-like manner. The ability of α-synuclein to cause multiple diseases is consistent with the strain hypothesis, which was introduced to explain the discrete neurological phenotypes induced by the prion agent.^1,2^ As research progressed, each prion strain became operationally defined as a particular phenotype observed under a fixed set of experimental parameters (reviewed in^3^). Consistent with these data, distinct sources of pathogenic α-synuclein exhibit discrete biological and biochemical properties in transmission models.^4–19^ However, as recent advances in cryo-electron microscopy (cryo-EM) enabled the determination of atomic resolution structures of misfolded proteins isolated from human and rodent samples, strain definitions are shifting away from one that is phenotypic toward one with a structural basis. For example, cryo-EM studies have now shown that while α-synuclein misfolds into amyloid filaments in patients with MSA,^20,21^ PD,^22^ and juvenile-onset synucleinopathy, the conformation of those filaments is distinct in each disease.^23^

This fundamental advance in our ability to interrogate the structural underpinnings of biological strains offers an unprecedented opportunity to reassess the systems we use to model disease and support preclinical testing in drug discovery programs. Synucleinopathies remain invariably fatal disorders, with countless therapeutic strategies failing to translate from bench to bedside. One hypothesis underlying these recurring setbacks is that the α-synuclein structures present in commonly used synucleinopathy models fail to recapitulate the α-synuclein folds present in human patients. Among these models, the TgM83 mouse line is commonly used to investigate disease pathogenesis and strain biology, in addition to preclinical evaluation of therapeutic candidates. These mice use the *Prnp* promoter to express human α-synuclein with the A53T mutation.^24^ TgM83^+/+^ mice develop spontaneous disease with hindbrain neuropathology in ~12-20 months, unlike TgM83^+/-^ animals, which remain asymptomatic over 2 years. However, following intracranial (i.c.) inoculation of pathogenic α-synuclein (reviewed in^25^), including human MSA patient samples, TgM83^+/-^ mice develop a terminal neurological disease.^4,10^ Biochemical analyses of α-synuclein in TgM83^+/+^ mice indicate these animals develop at least 3 distinct strains, all of which differ from human synucleinopathy samples.^16^ By comparison, TgM83^+/-^ mice develop clinical signs in ~4 months, and the resulting α-synuclein strains are consistent with the inoculum used in each experiment.^4–6,26^ Similar studies in mice expressing wild-type human α-synuclein require ~9.5 months for clinical onset.^10^ Because of the shortened timeframe and retention of strain properties, inoculation studies in the TgM83^+/-^ model have become pervasive throughout the synucleinopathy field. However, while samples from both spontaneously ill and inoculated TgM83 mice have been evaluated biochemically and biologically for maintenance of strain properties, very little is known about whether the resulting α-synuclein fibrils recapitulate the conformations found in human patient samples.^27,28^

Addressing this critical knowledge gap and empowering improved model development, here we report the first structural determination of α-synuclein fibrils from an MSA patient sample before and after transmission to TgM83^+/-^ mice. We show that key structural aspects of the MSA fibril fold are preserved upon mouse transmission, though notable and important differences indicate that MSA α-synuclein adapts to the new host during replication. Moreover, both the human and mouse-passaged MSA conformations are distinct from the heterogeneous population of α-synuclein fibrils that spontaneously form in TgM83^+/+^ mice. These findings demonstrate that by retaining strain (i.e., structural) properties during passage in TgM83^+/-^ mice, we validate the unique merits of this model for supporting preclinical research as well as confirm that MSA replicates using the prion mechanism of disease.

## RESULTS

### Resolution of α-synuclein fibrils from patient sample MSA14

To establish the structural basis for MSA passaging studies (Fig. 1a), we first determined the fold of α-synuclein filaments isolated from human patient sample MSA14 (used in previous studies^5,6,10,26,29^, Fig. 1b). Sarkosyl-insoluble filaments extracted from the anterior internal capsule were evaluated via negative stain and cryo-EM (Supplementary Fig. 1a; Supplementary Table 1). After data collection, image processing, and helical reconstruction, a single filament fold with two asymmetric protofilaments was resolved to 2.6 Å (Fig. 1c, Supplementary Fig. 1a-f, Supplementary Table 1). Despite extensive classification and the use of multiple initial models to search for low-population folds, only the one filament form was observed. The larger protofilament contains a characteristic cross-β hairpin and single-layered L-shaped motifs at one terminus and 3-layered L-shape at the other terminus, while the smaller protofilament is comprised of a cross-β hairpin and a U-shaped motif at the termini. There is an extended protofilament interface spanning one side of the hairpin- and L-motifs in PF-IA and a hammerhead-like region of PF-IB. Between the two protofilaments exists a pocket surrounded by basic residues, which contains additional density, likely a negatively charged small molecule or metabolite (Supplementary Fig. 1e-f). The ordered core consists of residues G14–F94 in the larger protofilament (PF-IA) and K21–Q99 in the smaller protofilament (PF-IB; Fig. 1c). This fold is almost identical to the published MSA Type I_1_ fold [C_α_ root mean square deviation (RMSD) = 0.503 Å; Supplementary Fig. 1g].^20^

**Fig. 1.**
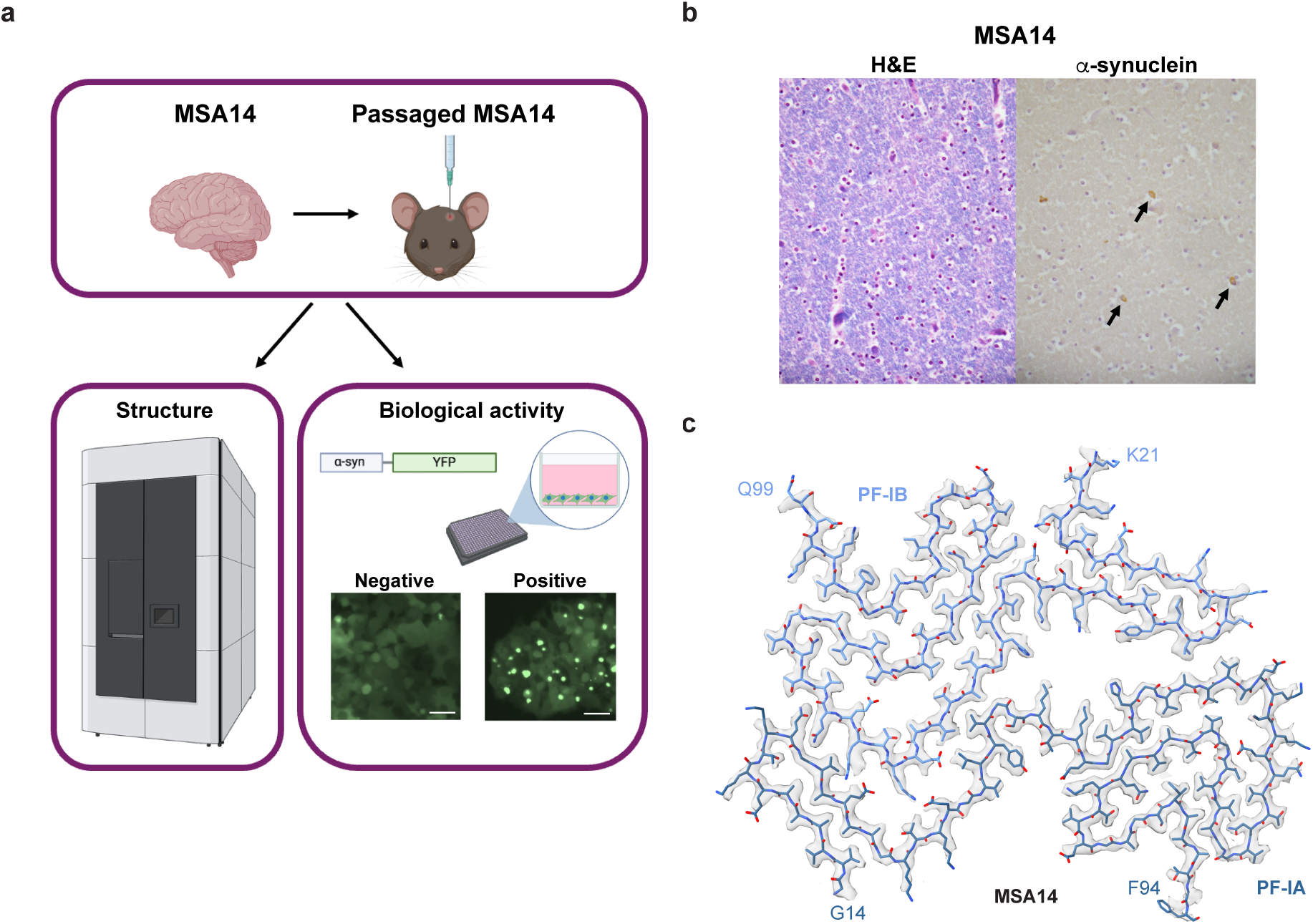
Resolution of the α-synuclein filament structure in patient sample MSA14. **a**, Schematic of experimental design. Homogenized tissue from the midbrain of patient sample MSA14 was inoculatd into TgM83^+/-^ mice. Sarkosyl-insoluble α-synuclein fibrils were isolated from both samples for structural (cryo-EM) and biological (biosensor cell assay) analysis. **b**, Representative neuropathology in the anterior limb of the internal capsule from patient sample MSA14, with hematoxylin and eosin on the left and immunostaining for α-synuclein on the right. **c**, Cryo-EM structure of α-synuclein filaments purified from the anterior internal capsule of patient MSA14. The atomic model (blue) fit into the cryo-EM density map (grey). This fold is consistent with Type I_1_ MSA filaments as previously reported.^20^

### MSA14 α-synuclein fibrils adapt to a new host during mouse transmission

Given our recent findings that α-synuclein strain properties in MSA patient samples appear to change, or adapt, after transmission to the TgM83^+/-^ mouse model,^11^ we sought to resolve the α-synuclein fibril structure of mouse-passaged MSA14 (Fig. 1a). TgM83^+/-^ mice were i.c. inoculated with either control patient sample C3 or MSA14 at 6-weeks-old. While the C3-inoculated mice remained asymptomatic through 365 days post-inoculation (dpi), MSA14-inoculated mice developed neurological disease in 201 ± 72 dpi (Fig. 2a; *P* = 0.0008). Brain samples from all C3-inoculated mice were collected with one hemisphere fixed for neuropathological analysis and one hemisphere frozen (n = 8). Five of the MSA14-inoculated mouse brains were similarly collected while the others were collected by microdissecting caudal brain regions only for cryo-EM structural determination. We quantified phosphorylated α-synuclein inclusions (pS129) in the hippocampus (HC), thalamus (Thal), hypothalamus (HTH), midbrain (Mid), and hindbrain (Hind) to confirm the caudal distribution of neuropathology (Fig. 2b; Supplementary Fig. 2). While neither inoculation cohort developed pathology in the HC (C3: 0.1 ± 0.1%; MSA14: 0.0 ± 0.0%; *P* = 0.9902) or Thal (C3: 0.1 ± 0.1%; MSA14: 0.0 ± 0.0%; *P* > 0.9999), MSA14-inoculated mice developed significantly more inclusions in the HTH (C3: 0.1 ± 0.0%; MSA14: 0.5 ± 0.3%; *P* = 0.0133), Mid (C3: 0.1 ± 0.0%; MSA14: 1.7 ± 0.3%; *P* < 0.0001), and Hind (C3: 0.0 ± 0.0%; MSA14: 1.9 ± 0.6%; *P* < 0.0001).

**Fig. 2.**
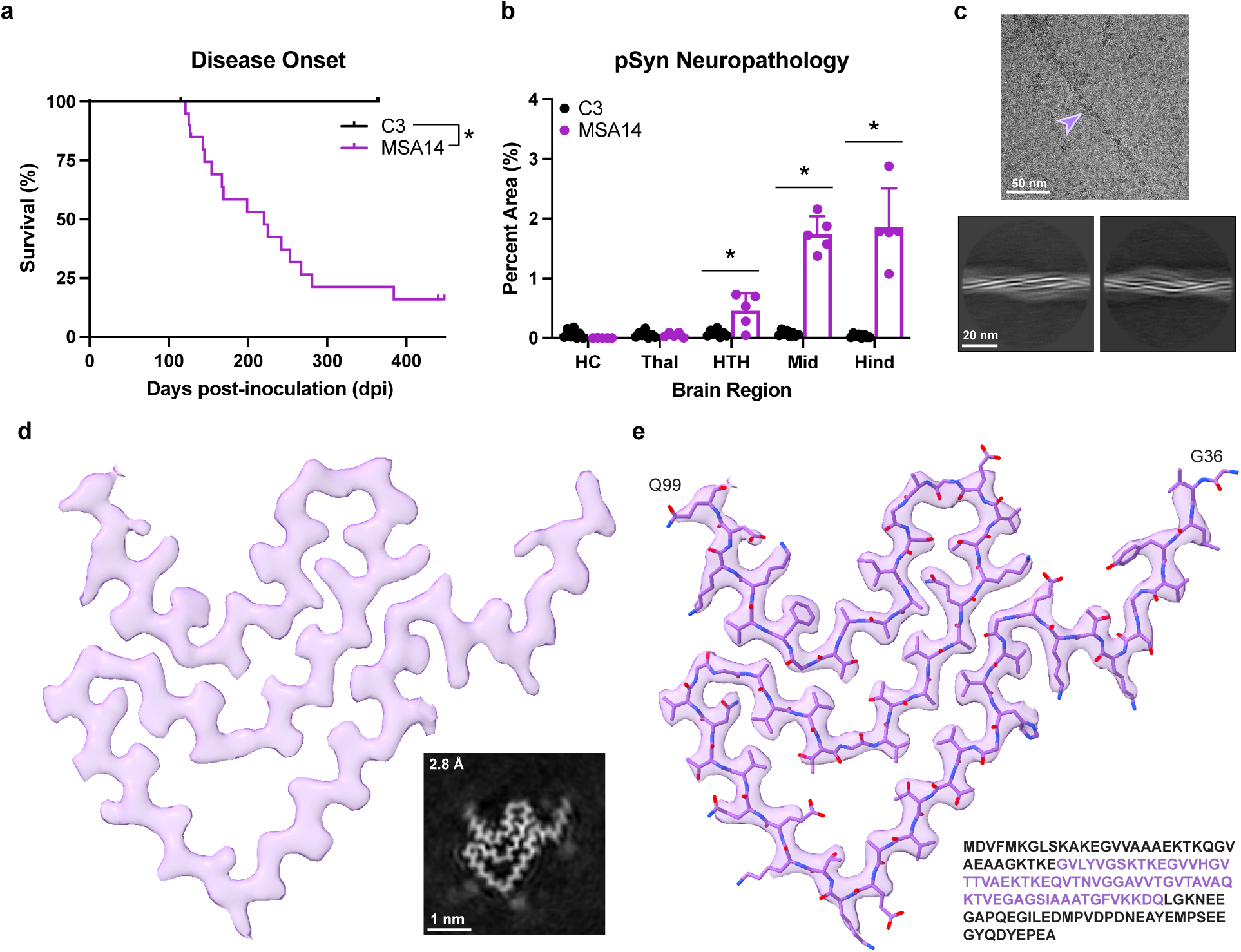
Adaptation of MSA α-synuclein fibrils to the TgM83^+/-^ host. **a**, Kaplan-Meier survival curves of TgM83^+/-^ mice intracranially inoculated with either patient sample C3 (black, n = 9) or MSA14 (purple, n = 18). **b**, Quantication of phosphorylated α-synuclein neuropathological inclusions in the hippocampus (HC), thalamus (Thal), hypothalamus (HTH), midbrain (Mid), and hindbrain (Hind) of TgM83^+/-^ mice inoculated with either C3 (n = 8) or MSA14 (n = 5) patient samples. **a & b**, *\* = P* < 0.05. **c**, Representative cryo-EM micrograph showing a TgM83^+/-^-passaged MSA14 α-synuclein fibril (purple arrow, top) and representative 2D class averages (bottom, 785 Å box), showing a morphology distinct from human MSA filaments. **d**, Cryo-EM map of mouse-passaged MSA14 fibrils, resolved to 2.8 Å, revealing a single protofilament whichpartially recapitulates PF-IB from the human MSA14 fold. **e**, Atomic model of α-synuclein residues 36-99 fit into the cryo-EM map from (**d**).

To investigate the structural fidelity of mouse-passaged MSA, we pooled two of the frozen TgM83^+/-^ hindbrains and purified sarkosyl-insoluble α-synuclein fibrils. The filaments were first evaluated via negative stain EM (Supplementary Fig. 3a), which showed clearly defined filaments with a similar width and length to those seen in the negative stain of the human patient (Supplementary Fig. 1a). However, the mouse-derived filaments appear slightly thinner with a more visible twist, and exhibit less linearity compared to the human-derived filaments.

After cryo-EM data collection, image processing, and helical reconstruction, we determined the cryo-EM structure of the mouse-passaged filaments to a resolution of 2.8 Å (Fig. 2c-e; Supplementary Figure 3; Supplementary Table 1), with clearly resolvable side chains and a rise of 4.75 Å and twist of −1.59°. Compared to the human filament, which has a rise of 4.76 Å and twist of −1.42°, the mouse-passaged filament exhibits a slightly greater twist, which agrees with a shorter crossover distance (525 Å) estimated from the 2D class averages (Fig. 2c). Notably, the density map shows only a single protofilament comprised of three layers, with a hammerhead motif opposite two grooves at the termini of the resolvable region (Fig. 2d). A U-shaped channel is present at one end of the ordered core, while an L-shaped motif at the other terminus forms the basis of the opposite groove. Modeling analysis revealed that residues 36-99 of α-synuclein make up the ordered core of the mouse-passaged protofilament (Fig. 2e). In comparison with the human MSA14 filaments inoculated into these mice, this protofilament structure matches closely with PF-IB (Fig. 3a). The human PF-IB and mouse-passaged folds are almost identical from residue T45 to the C-terminus (Q99), containing the same U-shaped motif at the C-terminus (A85-Q99) and a 2-layered hammerhead in the central residues (T45-A78, C_α_ RMSD = 0.606 Å). However, the characteristic hairpin (residues 21-35) of the N-terminal arm of PF-IB is absent, replaced by a shorter, single-layered L-shaped motif at the N-terminus (G36-E46) of the mouse-passaged MSA14 structure. Along with the N-terminal hairpin, the entirety of PF-IA is not resolvable in the ordered core of the mouse-passaged filaments. Comparing the structures down the filament axis (Fig. 3a, inset), the human Type I_1_ fold and the mouse-passaged MSA14 fold match well in the Z plane throughout the C-terminal body (G47-Q99) but diverge at the N-terminus (G36-E46), the same area where the folds differ in the cross-sectional view.

**Fig. 3.**
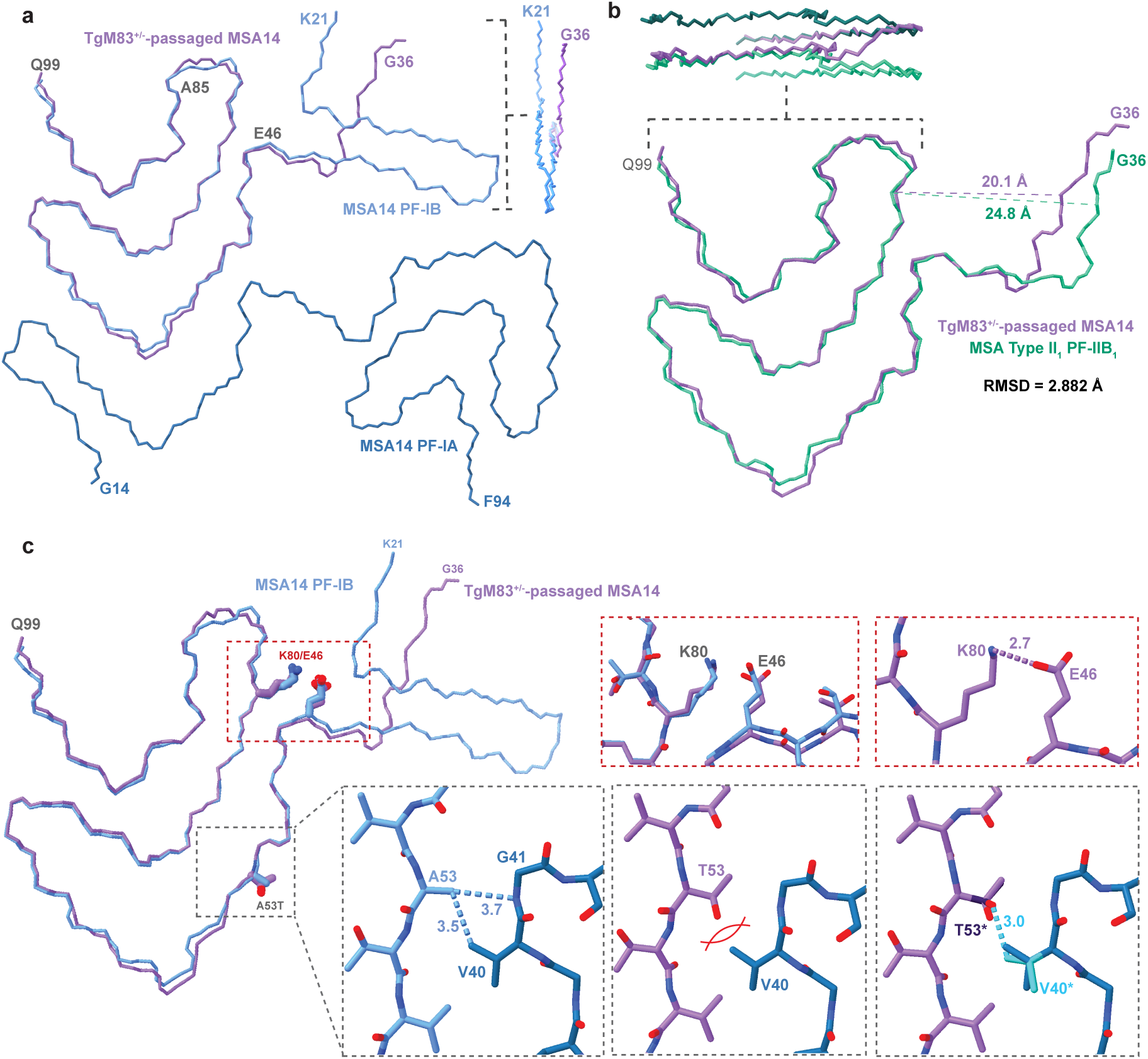
Mouse-passaged MSA14 fibrils retain core facets of the MSA14 fibril structure. **a**, Cross-section of model overlaying the MSA14 structure (blue) with the TgM83^+/-^-passaged MSA14 fibril (purple). **b**, Cross-sectional view of the model overlaying PF-IIB_1_ from the MSA Type II_1_ structure (green; PDB: 6XYP^20^) with the TgM83^+/-^-passaged MSA14 fibril (purple). Insets in (**a**) and (**b**) show differences between mouse-passaged and human filaments in the filament plane. **c**, Cross-section view of PF-IB from the MSA14 fibril structure (blue) overlaid with the TgM83^+/-^-passaged MSA14 structure (purple). Top insets (red) show the preserved salt bridge between E46 and K80. On the bottom, (left) the interaction in MSA14 fold between A53 and V40 of PF-1B and PF-1A, respectively, is incompatible (middle) with the A53T rotamer modeled in the mouse-passaged structure. (Right) However, slight rotations of the V40 (light blue) and T53 (dark purple) sidechains accomodates the interaction.

Interestingly, of all the human MSA folds resolved to date, the mouse-passaged fold most closely matches PF-IIB_1_ from the Type II_1_ human MSA structure (PDB: 6XYP; PF-IIB_1_ RMSD = 2.882 Å, PF-IB RMSD = 6.058 Å; Fig. 3b).^20^ As in PF-IB, residues 45-99 match the mouse-passaged fold (C_a_ RMSD = 0.743 Å), however, the N-terminal residues in PF-IIB_1_ are more closely aligned than those from PF-IB, with PF-IIB_1_ having the same short, single-layered L-shaped arm as the mouse-passaged fold. The groove formed by this arm and residues K80-D83 is narrower in the mouse-passaged fold (Fig. 3b), differentiating the two. A portion of the mouse-passaged fold is also Z-displaced from PF-IIB_1_, although here the difference arises at the C-terminal U-shaped motif. Starting at G86, the mouse-passaged fold protrudes by almost exactly one rung of the filament, closely matching the adjacent strand of PF-IIB_1_ in both the Z-plane and cross-section (Fig. 3b, top). The relatively small, flexible residues in this region (G84, A85, G86) may impart unique degrees of freedom on this area of the filament, as there is also a deviation between the mouse-passaged MSA14 fold and PF-IB at these residues, despite the chains aligning almost perfectly on either side of this segment.

### Key structural facets of MSA14 fibrils are maintained upon mouse passage

Notably, the TgM83 mouse model expresses the Parkinson’s-causing A53T mutation, though no α-synuclein mutations have been identified in an MSA patient to date. In all human MSA folds, A53 in PF-B lies at the protofilament interface and makes van der Waals contacts (3.5-3.7 Å) with V40 in PF-A (Fig. 3c, bottom left inset, shown in the MSA14 structure). In our mouse-passaged MSA14 structure, T53 lies unperturbed relative to A53 in the human fold, with well-defined density for the sidechain (Supplementary Fig. 4a). When replacing PF-IB in our human structure with the mouse-passaged structure, T53 clashes with the sidechain of V40 in PF-IA (O-C distance 2.3 Å; Fig. 3c, bottom middle inset). However, a slight rotation (10-20°) of each sidechain restores the steric tolerance for T53 at the protofilament interface (Fig. 3c, bottom right inset), with no intra-protofilament clashes on adjacent rungs (Supplementary Fig. 4a, right inset). Furthermore, A53 in PF-A makes cross-protofilament contacts with either V37 (MSA Type I) or V40 (MSA Type II), and a threonine substitution is again tolerated (Supplementary Fig. 4b & c). As these are minor sidechain rearrangements, we believe that this substitution has limited influence on the structural differences between the human and mouse-passaged MSA14 folds, and that the increased propensity of A53T to propagate MSA α-synuclein^5,6,10^ may arise from other factors. This effect was clearly seen when we tested both MSA14 and mouse-passaged MSA14 samples in our panel of α-syn-YFP biosensor cell lines (Table 1). In comparison with other mutations, both the human and mouse-passaged sample showed the greatest propensity to replicate in the α-syn*A53T-YFP cell line.

**Table 1.** A-synuclein biosensor cell assay data.

| Mutation | Human Patient Samples<sup>a</sup> |  |  | Mouse-Passaged Samples |  |  |
| --- | --- | --- | --- | --- | --- | --- |
|  | C3 | MSA14 | *P* Value | C3<sup>b</sup> | MSA14<sup>c</sup> | *P* Value |
| A30G | 0.5 ± 0.6 | 3.8 ± 2.0 | 0.0002 | 1.0 ± 0.4 | 3.5 ± 1.7 | 0.0306 |
| E46K | 0.5 ± 0.3 | 1.0 ± 0.5 | 0.0501 | 0.6 ± 0.3 | 0.6 ± 0.4 | 0.9999 |
| G51D | 0.7 ± 0.6 | 2.1 ± 1.4 | 0.0675 | 1.2 ± 0.4 | 1.9 ± 0.6 | 0.0485 |
| A53E | 3.7 ± 1.1 | 1.7 ± 1.1 | 0.0093 | 1.2 ± 0.2 | 1.6 ± 0.5 | 0.1964 |
| A53T | 3.1 ± 1.0 | 23 ± 11 | 0.0075 | 3.6 ± 0.5 | 16 ± 7.5 | 0.0207 |
| V55Y | 1.0 ± 0.4 | 3.5 ± 0.5 | 0.0029 | 1.6 ± 0.5 | 4.8 ± 0.8 | 0.0033 |
| V66F | 1.5 ± 0.6 | 2.0 ± 1.4 | 0.4711 | 2.8 ± 1.2 | 3.1 ± 0.8 | 0.5237 |
| V74P | 0.4 ± 0.2 | 0.9 ± 0.4 | 0.0284 | 0.4 ± 0.2 | 0.8 ± 0.6 | 0.1753 |

Data reported as average ± SD. <sup>a</sup>Data represent n = 6 technical replicate wells. <sup>b</sup>Data represent n = 8 biological replicates. <sup>c</sup>Data represent n = 5 biological replicates.

In contrast to A53T, the PD-causing E46K mutation blocks propagation of MSA^6,9^ but facilitates replication of α-synuclein from patients with Lewy body diseases (PD, dementia with Lewy bodies, or Parkinson’s diseases with dementia).^6,8^ Cryo-EM studies have shown that the structural basis for this phenomenon is due to the presence of a stabilizing salt bridge formed between E46 and K80 in the B protofilaments of all human MSA folds, and the A protofilaments of Types I_1_, II_1_ and II_2_ filaments, while E46 faces solvent in the α-synuclein fold found in patients with the Lewy body diseases (the Lewy fold).^20–22^ Mutations to either E46 or K80 introduce repulsive forces that prevent MSA propagation in cellular models.^30^ Consistent with this defining feature of the MSA fold, the mouse-passaged MSA14 fibril also maintained the E46/K80 salt bridge (Fig. 3c, top insets). Moreover, when we tested both MSA14 and mouse-passaged MSA14 samples in the α-syn*E46K-YFP cell line, none of the samples were able to replicate (Table 1).

Apart from these two mutations, we evaluated the infectivity profile of MSA14 relative to mouse-passaged MSA14 in additional mutant cell lines (Table 1). Both human and mouse-passaged MSA14 exhibited mild to weak replication in the A30G and V55Y cells, but were unable to propagate using G51D, A53E, V66F, or V74P mutant protein. Analyzing these results through a structural lens, the decreased or inability of V55Y and V74P to propagate is unsurprising, given the productive interactions made by the valine sidechains in a sterically constrained environment. The same holds true for A53E in cells incubated with the MSA14 sample. An aspartate residue at position 51 may clash with the sidechain of H50, and the decrease in flexibility caused by this substitution could also make a turn, which is seen in both human and mouse folds at this position, unfavorable. However, the inability of the mouse-passaged samples to propagate using A53E α-synuclein, as well as the lack of replication of human and mouse samples using V66F α-synuclein, are more surprising considering that these residues face solvent and both steric and electrostatic alterations should be tolerated. A30G should also be tolerated in both A and B human protofilaments and the mouse-passaged structure, making the decrease in infectivity in this cell line unexpected as well.

### Endogenous α-synuclein is not incorporated into mouse-passaged MSA14 fibrils

As the TgM83^+/-^ mice used here also express endogenous mouse α-synuclein, we sought to determine whether the mouse-passaged filaments are comprised of mouse α-synuclein, human α-synuclein, or a mixture of both. While there are seven amino acid differences between the two species, only two are in the ordered region of the filament: residue 53 (alanine in human and threonine in mouse) and residue 87 (serine in human and asparagine in mouse). Given that TgM83^+/-^ mice already express the A53T mutation, the only sequence difference in the resolved structure is S87N. In the mouse-passaged MSA structure, S87 faces inward toward a sterically constrained pocket where it engages in hydrogen bonding with T81 (Fig. 4a inset). When an asparagine is directly substituted into position 87 of the model, the residue protrudes out of the well-defined side-chain density and clashes with T81 on the opposite side of the hairpin loop with only 1.9 Å separation (Fig. 4b). An altered N87 rotamer allows for the for a conformation with relatively short interatom distances (2.8-2.9 Å; Fig. 4c), although these distances have been observed previously in other constrained systems.^31^ However, given that this asparagine orientation does not match the well-defined sidechain density, it is highly likely that only human α-synuclein accumulates into these filaments.

**Fig. 4.**
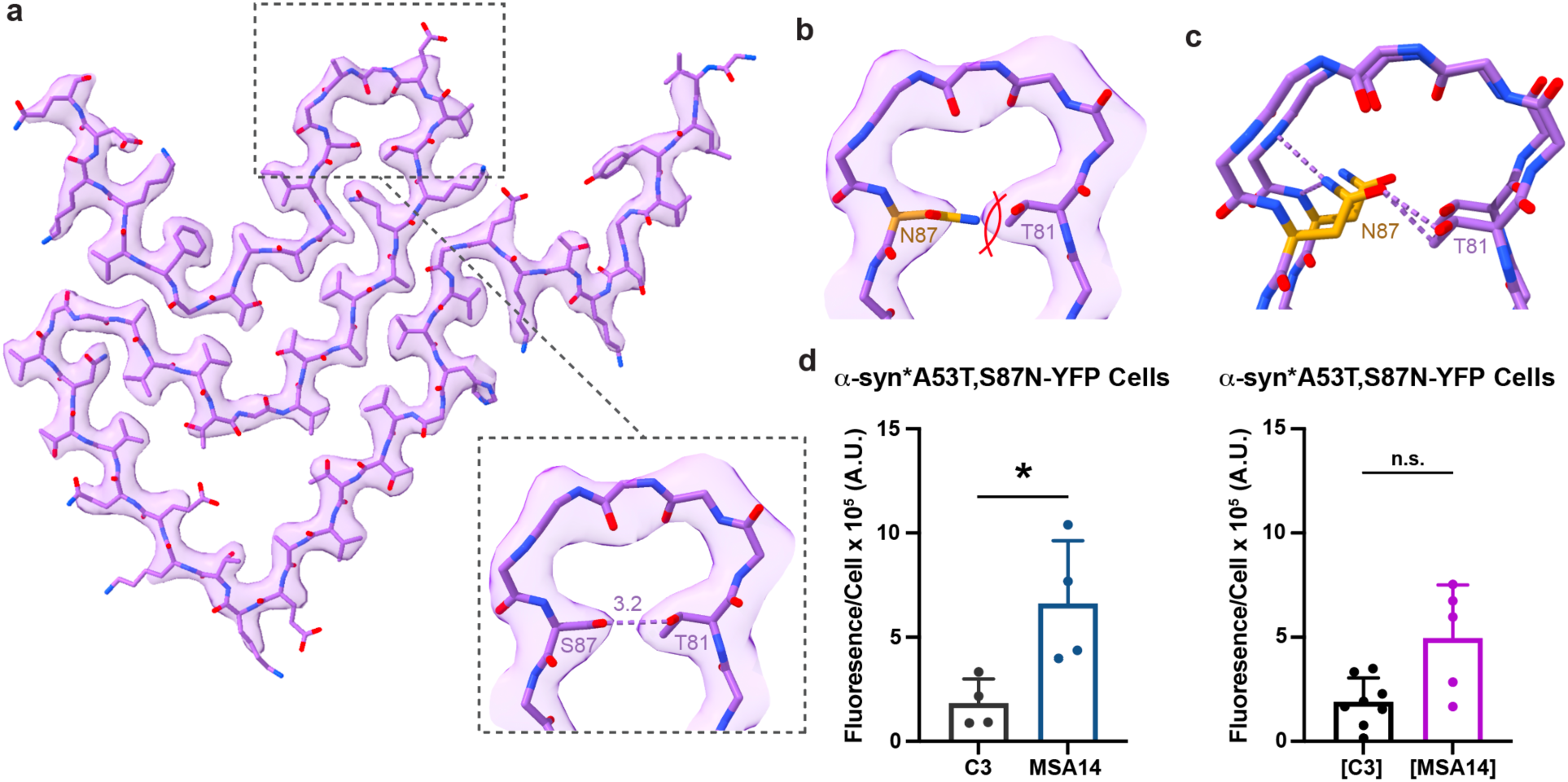
Mouse-passaged MSA14 selectively replicate using human α-synuclein *in vivo*. **a**, Density map from TgM83^+/-^-passaged MSA14 fibrils is consistent with the human serine at position 87 hydrogen bonding with T81. **b,** The murine synuclein S87N mutation does not fit the cryo-EM density, and clashes with T81. However, a different rotamer (**c**) is accomodated, with 2.8-2.9 Å distances between the asparagine sidechain and surrounding atoms in the pocket. **d**, Cell infection data comparing (left) C3 and MSA14 patient samples and (right) mouse-passaged C3 ([C3]) and MSA14 ([MSA14]) on the α-syn*A53T,S87N-YFPcells [x 10^5^ Arbitrary Units (A.U.)]. * = *P* < 0.05.

To determine the ability of the MSA14 and mouse-passaged MSA14 fibrils to replicate using α-synuclein containing the mouse substitutions, we generated monoclonal HEK293T cells expressing α-syn-YFP with the A53T and S87N mutations. MSA14 showed a significant ability to replicate in the cells (Fig. 4d; 6.6 ± 3.0 x 10^5^ A.U.; *P* = 0.0436) compared to C3 (1.8 ± 1.2 x 10^5^ A.U.). Interestingly, while mouse-passaged C3 samples (indicated as [C3]) also showed no infection (1.9 ± 1.2 x 10^5^ A.U.), we observed variability with the mouse-passaged MSA14 samples (designated [MSA14]; 5.0 ± 2.6 x 10^5^ A.U.). Some of the samples successfully propagated in the α-syn*A53T,S87N-YFP cells whereas others did not, resulting in an insignificant difference compared to [C3] (*P* = 0.0530). These data are consistent with our structural findings that the MSA14 fibrils prefer to replicate using S87 α-synuclein, but both are able to adapt to the change in primary sequence when N87 α-synuclein is the only substrate available.

### The mouse-passaged MSA14 fold includes additional densities

Strikingly, while the mouse-passaged filament fold closely resembles PF-IB of the human inocula, there is no second protofilament resolvable in the mouse structure, as is the case in all human MSA folds. During 3D classification, we were able to resolve two lower-resolution maps of the same overall mouse-passaged fold, but which contain three obvious additional densities protruding from the filament backbone (Fig. 5a & b; Supplementary Fig. 3a). Examining the highest resolution mouse-passaged filament map (Fig. 2d) low-pass filtered to 7.5 Å (σ = 3.25) also reveals these densities, extending from the sidechains of K43, K58, and K60 (Fig. 5c). These each appear connected to the terminus of the lysine sidechains, extending approximately 20 Å, and are roughly perpendicular to the filament backbone. As such, we hypothesize that these are post-translational modifications to the lysine residues, or non-covalent densities bound at the exterior of the filament. In the human MSA14 fold, the N-terminal hairpin clashes with the density extending from K43, via the sidechain of Y39 (Fig. 5d). K43 also points inward toward the unidentified, likely negatively charged, density between the two protofilaments of the human MSA fold (Supplementary Fig. 1f) and may destabilize heterodimer formation. Furthermore, the densities associated with K58 and K60 in the mouse-passaged maps would interfere with productive cross-protofilament interactions in the human fold, where K58 and K60 of PF-B form salt bridges with E35 and E28 of PF-A in the human fold, respectively (Fig. 5e & f). Overall, we hypothesize that these extra entities preclude the attachment of a second protofilament and the formation the N-terminal hairpin in PF-IB, and as such are responsible for incomplete recapitulation of the human MSA14 fold.

**Fig. 5.**
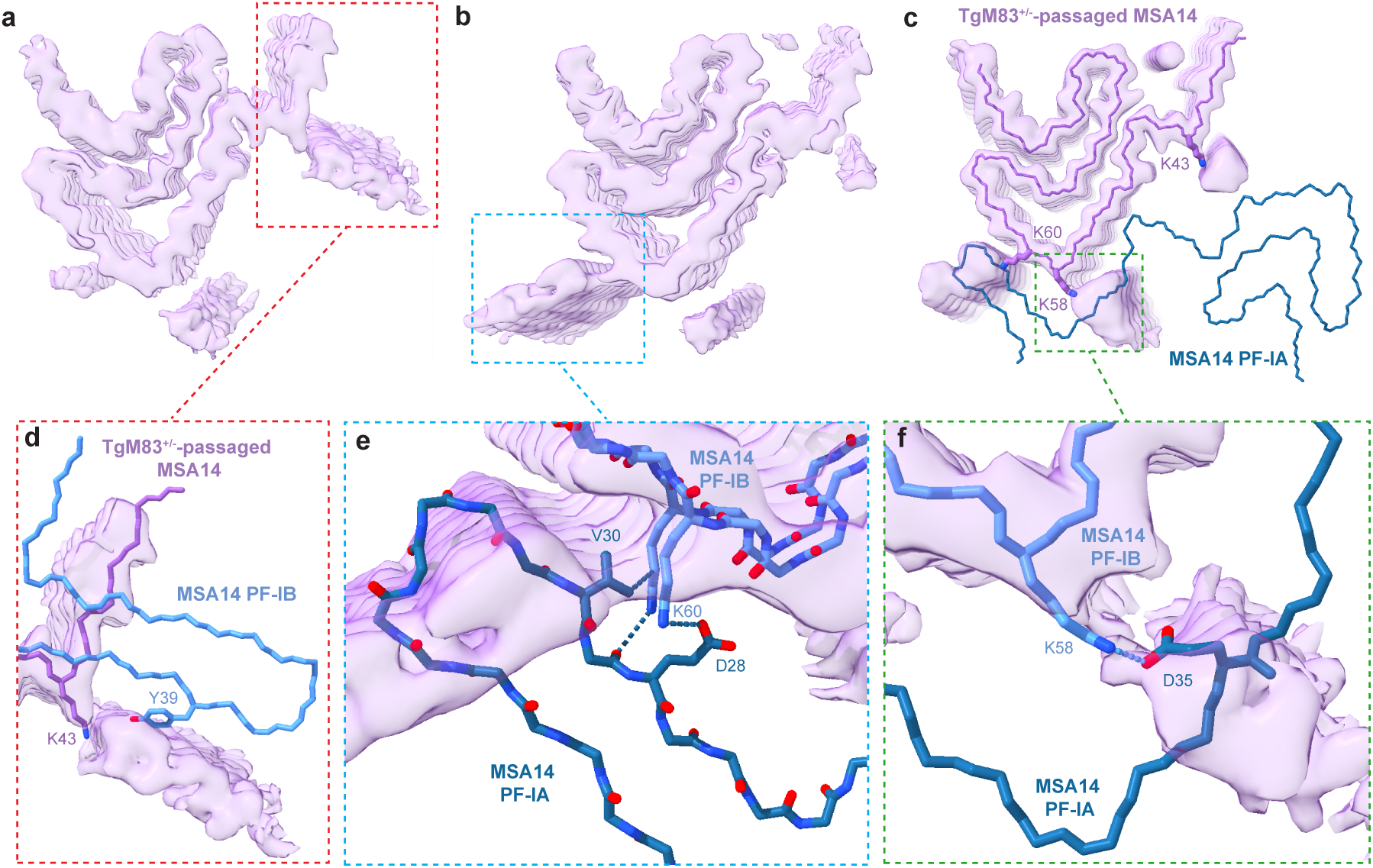
Additional densities on mouse-passaged MSA14 fibril preclude protofilament formation. **a & b**, Maps with additional densities resolved during 3D classification, which are also observed in the canonical map (**c**) low-pass filtered to 7.5 Å (σ = 3.25) and appear to extend from the ends of the sidechains of K43, K58, and K60. Overlaying the mouse-passaged structure purple onto PF-IB of the MSA14 structure, these densities are incompatible with the presence of PF-IA (blue). **d**, The density extending from K43 in the mouse-passaged structure is incompatible with Y39 in the MSA14 structure. **e & f**, Overlay of the mouse-passaged MSA14 density (purple) with the human MSA14 structure. **e**, Additional density extending from K60 precludes interactions in the human structure between K60 in PF-IB and V30 and D28 in PF-IA. **f**, The cross-protofilament salt bridge between K58 and D35 in the human structure is occluded by the additional density in the mouse passaged structure.

### TgM83^+/+^ mice spontaneously develop multiple α-synuclein fibril conformations

The TgM83^+/-^ mouse model is regularly used for inoculation studies as the mice do not develop spontaneous disease. By comparison, the TgM83^+/+^ model, which develops clinical disease in ~12-20 months,^16,24^ is used to investigate spontaneous synucleinopathy. To compare α-synuclein strain biology between MSA-inoculated TgM83^+/-^ mice with spontaneously symptomatic TgM83^+/+^ animals, we aged our retired male TgM83^+/+^ breeders until onset of progressive neurological signs. Brains from 6 terminal animals were collected half-fixed and half-frozen, 1 was fully frozen, and an additional 2 brains were collected by microdissecting the caudal brain regions for cryo-EM. As negative controls, we aged 2 retired B6C3F1 female breeders to 1 year and collected brains half-fixed and half-frozen. We first tested homogenized half-brains for α-synuclein prion activity in our α-syn*A53T-YFP cell assay (Fig. 6a). Despite all TgM83^+/+^ mice showing clinical signs at the time of collection, only one sample (M5) propagated in the biosensor cells (*P* < 0.0001). By comparison, mouse M2, which did not replicate in the α-syn*A53T-YFP cells, had robust α-synuclein neuropathology, most notably in the Mid. While mouse M5 also had Mid pathology, the total density was to a much lower extent, again demonstrating strain differences between the animals (Fig. 6b). So *et al*. recently reported three sub-type classifications of the TgM83^+/+^ strain based on proteinase K (PK) and thermolysin (TL) digestion patterns.^16^ Using the same digestion protocols and primary antibody, we observed some overlapping digestion patterns as previously reported, along with additional banding patterns (Fig. 6c & d; Supplementary Figs. 5 & 6). Moreover, while samples M5 and M6 were almost identical by protease digestion, the two samples differed substantially based on biological activity in the α-syn*A53T-YFP cell assay, suggesting that multiple α-synuclein fibril conformations develop in the mouse model.

**Fig. 6.**
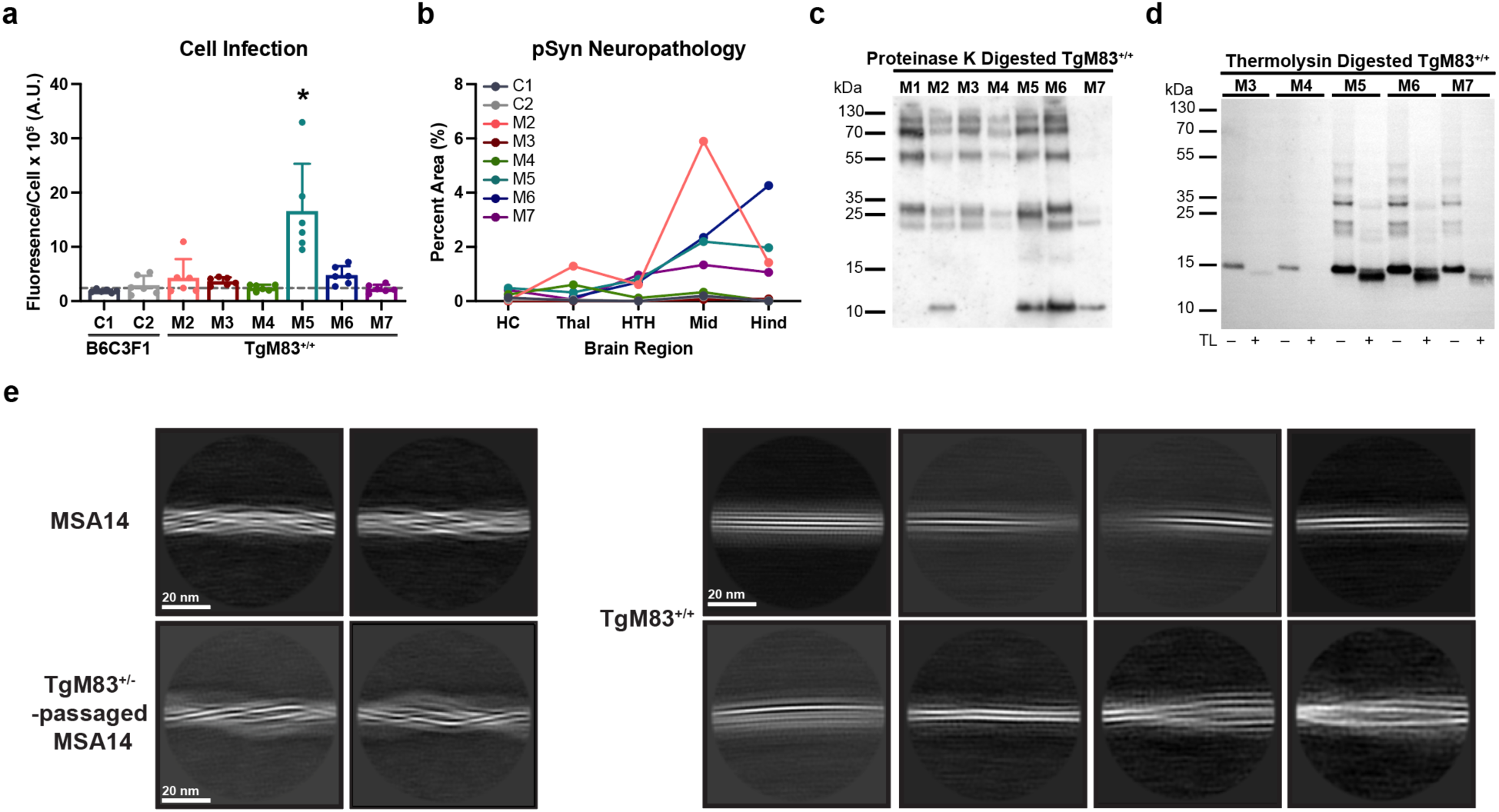
Spontaneously ill TgM83^+/+^ mice develop multiple α-synuclein fibril conformations. **a**, Cell infection data comparing non-transgenic B6C3F1 controls (in grays) with spontaneously symptomatic TgM83^+/+^ samples (in colors) measuring replication in α-syn*A53T-YFP cells [x 10^5^ Arbitrary Units (A.U.)]. Dashed gray line at 2.4 x 10^5^ A.U. indicates average value from negative control samples. * = *P* < 0.05. **b**, Quantication of phosphorylated α-synuclein neuropathological inclusions in the hippocampus (HC), thalamus (Thal), hypothalamus (HTH), midbrain (Mid), and hindbrain (Hind) of B6C3F1 controls or spontaneously ill TgM83^+/+^ mice. **c & d**, Western blot of (**c**) proteinase K (PK)-digested or (**d**) thermolysin (TL) digested insoluble α-synuclein isolated from spontaneously symptomatic TgM83^+/+^ mice. **c**, Primary antibody: Clone 42 (1:10,000). **d**, Primary antibody: EP1536Y (1:10,000). **e**, Representative 2D class averages of MSA14 fibrils (top left) and TgM83^+/−^-passaged MSA14 fibrils (bottom left). (Right) representative 2D class averages of α-synuclein fibrils from two pooled hindbrains of spontaneously symptomatic TgM83^+/+^ mice show heterogeneous filament morphologies, which do not resemble those from MSA14 or mouse-passaged MSA14.

In seeking to resolve the α-synuclein fold in TgM83^+/+^ mice, we pooled hindbrains dissected from two terminal animals and isolated sarkosyl-insoluble fibrils. Negative stain and cryo-EM analysis indicated a heterogenous mixture of filament types. The differences in these filament classes are identifiable at the micrograph scale as well as in 2D classification, which highlighted at least 5 distinct subpopulations (Fig. 6e). It is clear from visual inspection that the class averages from the TgM83^+/+^ data do not resemble those from either the MSA14 or mouse-passaged MSA14 datasets. Therefore, it is likely that these filaments adopt entirely different folds from those seen in humans. Indeed, recently two TgM83^+/+^ filament folds, which contained a mixture of human and mouse α-synuclein, were solved to high-resolution (the mA53T fold).^28^ The large majority of these filaments were a single, 3-layered protofilament comprised of residues 36-99 with an L-shaped motif at the N-terminus, matching our mouse-passaged structure. However, the mA53T structure has a more compact fold, with extended packing interfaces compared to the mouse-passaged MSA14 structure, as well as a more open C-terminal channel notably able to accommodate S87N, which faces solvent. Neither this singlet fold, nor the less abundant doublet of the same protofilament, recapitulated any of the human MSA folds, or any other disease-relevant α-synuclein structure, leading to the conclusion that TgM83^+/+^ mice are a poor model for human synucleinopathies.

## DISCUSSION

The ability of MSA patient samples to transmit neurological disease to TgM83^+/-^mice was first reported using tissue from two patients in 2013.^4^ In the following years, through the use of animal models and corresponding cell-based assays, increasing evidence has continued to support the classification of MSA as a prion disease.^5,7,10,11,13,26,32–34^ Over that same time, cryo-EM has proven a powerful tool for studying MSA and other neurodegenerative diseases by elucidating the structures of misfolded proteins in post-mortem patient tissue. However, in the absence of structural data from experimental transmission studies, questions remain about whether MSA animal and cellular models recapitulate disease relevant folds, or if this approach just accelerates spontaneous α-synuclein misfolding. Here, we report the first cryo-EM structures of any misfolded protein from both the starting human patient sample inoculum and the terminal mice following disease transmission. We find that the mouse-passaged MSA filaments consist of a single protofilament, which largely matches the fold of one protofilament found in the heterodimeric patient fibrils. Our data showing that pathogenic α-synuclein retains its fibril structure when replicating in TgM83^+/-^ mice aligns with recent reports from others investigating transmission of recombinant fibrils.^27,35^ However, here we find that the N-terminus of the replicated MSA protofilament and the entirety of the opposing protofilament are absent. While this indicates that MSA fibrils adapt to a new host, the fact that critical facets of the MSA α-synuclein fold are maintained during passaging demonstrates that the inoculated TgM83^+/-^ mouse can serve as a valuable model for understanding disease biology and supporting drug discovery programs. Moreover, additional densities extending from the main chain of the mouse-passaged MSA fold provide insight into the incomplete replication of the human MSA conformation upon transmission. Finally, given the heterogeneous fibril population present in spontaneously ill TgM83^+/+^ mice, the presence of a single α-synuclein fold containing only human α-synuclein in MSA-inoculated TgM83^+/-^ mice confirms that MSA replicates *in vivo* using the prion mechanism of disease.

Given our recent cell-based findings indicating that MSA strain properties are altered following transmission to the TgM83^+/-^ mouse model,^11^ we hypothesized that the observed adaptation was likely due to the presence of the PD-causing A53T mutation in the human α-synuclein transgene. As discussed above, this mutation sits in a sterically constrained area at the protofilament interface in the reported human MSA folds.^20,36^ Notably, MSA transmission studies using the TgM83^+/-^ mouse model are largely criticized, given that the A53T mutation is only present in familial PD cases and is not associated with MSA. It is, therefore, quite notable that our data indicate that the A53T mutation has limited, if any, impact on MSA14 adaptation to the TgM83^+/-^ host, as a slight rotation of the sidechains of T53 and V40 should accommodate the change in primary sequence. These data support the continued use of the A53T mutation in TgM83^+/-^ mice to accelerate α-synuclein replication, and subsequently MSA research, compared to the prolonged incubation periods seen in the wild-type expressing TgM20 model. Instead of the A53T mutation driving structural differences, we find that the presence of three large additional densities extending from lysine sidechains in the mouse-passaged structure may preclude full recapitulation of the MSA14 conformation, as they are incompatible with formation of the protofilament heterodimer and N-terminal hairpin seen in MSA14 filaments (Fig. 5). Two resides with additional densities, K58 and K60, form important contacts at the extended protofilament interface in all resolved human MSA structures. These densities, which are ~20 Å long and ~7 Å wide, would interfere with adjacent interface contacts as well, precluding the presence of a second protofilament in the same relative orientation as those found in human MSA structures. Furthermore, the density appearing from the sidechain of K43 also extends into the pocket where the non-proteinaceous cofactor seen in all MSA folds binds. This cofactor likely serves to satisfy the positively charged residues that point inward toward the binding pocket and contributes to the stabilization of the disease-relevant heterodimers. While it remains unknown if this cofactor is necessary for the second protofilament to bind (e.g., serves as a means for facilitating α-synuclein misfolding) or if it simply serves to mitigate repulsive positive charges between a previously formed heterodimeric structure, the inability of the cofactor to bind combined with the disruption of the extended protofilament interface likely plays a key role in a single filament fold forming in MSA-inoculated TgM83^+/-^ mice.

Unexpectedly, this finding provides intriguing insights into α-synuclein fibril replication in MSA. From our structural characterization here, PF-IB is a stable α-synuclein fold (i.e., PF-IB does not require PF-IA for formation nor is PF-IA needed to prevent degradation of PF-IB). While our studies are unable to determine when addition of the densities at the protofilament interface occurs during the replication process, the absence of PF-IA suggests a few potential underlying mechanisms. In the first, PF-IB forms first and is required for the formation of PF-IA. However, densities attach to K43, K58, and K60 imminently after fibril formation, interfering with subsequent templating of PF-IA. In the second, templating of PF-IB and PF-IA occurs independently, but PF-IA is either less stable in the TgM83^+/-^ mouse model and is subsequently degraded prior to protofilament formation, or it is temporally offset from PF-IB and has not formed yet. Then, in the absence of PF-IA attachment, the lysine residues are accessible for density binding. Regardless of which scenario occurs, PF-IA is does not attach to PF-IB, preventing the full recapitulation of the heterotypic MSA fold.

These results also indicate that MSA fibril replication selectively recruits human α-synuclein monomer, excluding mouse α-synuclein. While our cell assay data indicate that replication can occur using α-synuclein containing an asparagine as position 87, the cryo-EM density points to a preference for the human serine residue. These findings are not only consistent with prior studies investigating the human/mouse species barrier for α-synuclein,^37^ but they are also in agreement with known species barriers impacting transmission of prion diseases. For example, the hamster prion strain 263K is unable to replicate using mouse prion protein (PrP) due to an asparagine to tyrosine polymorphism at position 155/154, respectively, as Y155 creates steric hinderance with E152 in the 263K fold.^38^ As a result, inoculation of hamster prions into mice expressing both mouse and hamster PrP results in selective recruitment of hamster PrP during replication.^39^

The presence of human α-synuclein only in the mouse-passaged fibrils differs from recent studies reported by Zhang *et al.*, who resolved cryo-EM structures of α-synuclein filaments formed spontaneously in TgM83^+/+^ mice (mA53T fold), which contain both human and mouse α-synuclein.^28^ The authors reported a single protofilament structure making up 93% of their sample population along with a homotypic doublet comprising the other 7%, neither of which matches the MSA fold or known folds of other synucleinopathies. Like our mouse-passaged structure, the TgM83^+/+^ protofilament ordered core also contains residues 36-99 of α-synuclein, with a matching single layer L-motif and conservation of the E46/K80 salt bridge near the N-terminus. However, K58 is flipped inward in the homozygous mouse fold, creating a tighter cross-ß pocket in the hammerhead region than is found in the MSA or matching mouse-passaged folds. Importantly, residue 87 faces solvent, allowing the structure to accommodate both the human serine and mouse asparagine, which are both found in the mixed α-synuclein species in these filaments.

Apart from these high-resolution structural differences between the mA53T and mouse-passaged MSA folds, there also appears to be biological and biochemical differences between our cohort of TgM83^+/+^ mice and those used to resolve the mA53T fold. For example, the 2D class averages of our cohort (Fig. 6) suggest the presence of at least 5 different filament types, including but not limited to those seen in particles contributing to the mA53T folds. This heterogeneity is seen both via sample infectivity in the α-syn*A53T-YFP cell lines and the presence of α-synuclein neuropathological inclusions in terminal mice. While brains from clinically affected mice contained detergent-insoluble α-synuclein, only one animal (M5) developed a strain capable of replicating in our biosensor cell line (Fig. 6a). This is even more notable given the sequence homology between the *in vivo* and *in vitro* models. Importantly, this outcome is not due to difference in pathology load in the brain, as mice M2 and M6 contained a higher burden of α-synuclein inclusions, and mouse M7 had similar lesions to mouse M5 (Fig. 6b). Biochemically, all mouse samples tested contained PK- and TL-resistant protein, but the banding patterns were variable across the samples, consistent with underlying conformational differences in the α-synuclein fibrils (Fig. 6c & d; Supplementary Figs. 5 & 6). These data agree with recent work from So *et al*., identifying three distinct α-synuclein populations in TgM83^+/+^ mice.^16^ However, given the presence of additional banding patterns here and the detection of at least 5 fibril populations in our 2D classifications, we expect that additional conformations beyond those initial 3 are present in the mice. Altogether, these data indicate that spontaneous α-synuclein filament formation in TgM83^+/+^ mice is a stochastic process across mice and cohorts, leading to a pool of heterogeneous folds.

The biological and now structural differences between MSA-inoculated TgM83^+/-^fibrils and the spontaneously formed TgM83^+/+^ filaments raise important concerns about the use of the TgM83^+/+^ model to support preclinical development of therapeutics for synucleinopathy patients. Prior efforts to develop small molecule therapeutics for prion disease patients resulted in the identification of the 2-aminothiozole IND24,^40^ which can extend survival in prion-inoculated mice but is ineffective against human prion strains.^41^ Combining the growing biological, biochemical, and structural differences between the TgM83^+/+^ strain and patient-derived strains reported here and by others,^16,28^ with the now established existence of multiple α-synuclein fibril conformations that exist both between^16^ and within animals, there is substantial data indicating that using the TgM83^+/+^ model in preclinical studies is unlikely to support successful translation of therapeutics into the clinic. Instead, while our findings indicate that mouse passaging results in adaptation of the MSA14 conformation, they also suggest that the TgM83^+/-^inoculation model is a useful one for biological understanding and drug discovery in the MSA field. Potential modifications to this model, such as mutation of the lysine residues containing additional densities, may improve maintenance of strain properties and structural fidelity.

## CONCLUSION

In resolving the cryo-EM structure of α-synuclein filaments from an MSA patient sample before and after transmission to the TgM83^+/-^ mouse model, we show that key structural aspects of PF-IB are maintained as the MSA patient sample adapts to a new host. For example, the resulting fibril conformation is nearly identical to that of the human inoculum for a majority of the passaged filament fold and can accommodate the A53T mutation by slightly rotating sidechains at positions T53 and V40. Additionally, the defining E46/K80 salt bridge is maintained, resulting in an inhibitory effect of the E46K mutation on *in vitro* replication. Importantly, the density map from the mouse-passaged MSA fold is incompatible with the endogenous mouse α-synuclein sequence, demonstrating that MSA replicates via selective recruitment of templating-competent monomer. We also identified large densities on three lysine residues in PF-IB, two of which preclude heterodimerization of the full MSA fold. Finally, while we find that MSA inoculation into TgM83^+/-^ mice induces replication of the MSA fold via the prion mechanism, stochastic misfolding of α-synuclein in the TgM83^+/+^ model results in a pool of heterogeneous folds, none of which are consistent with patient-derived structures. This structural variability, paired with inconsistent biological and biochemical outcomes, indicates that TgM83^+/+^ mice should not be used as a model of human synucleinopathy.

## MATERIALS AND METHODS

### Data availability

Cryo-EM maps and atomic coordinates are deposited in the EMDB and PDB with access codes: XXXXX and YYYY. All other data generated or analyzed during this study are included in this published article and accompanying supplementary information files.

### Human tissue sample and immunostaining

The Massachusetts Alzheimer’s Disease Research Center provided frozen brain tissue samples from a patient with neuropathologically confirmed MSA (MSA14). The fresh brain was bisected longitudinally with one half coronally sectioned and rapidly frozen while the other half was fixed in 10% (wt/vol) neutral buffered formalin prior to sectioning. Neuropathological evaluation was done on a set of blocked regions representative for a variety of neurodegenerative diseases. All blocks were stained with Luxol fast blue and hematoxylin and eosin. Selected blocks were used for immunostaining with α-synuclein, ß-amyloid, and phosphorylated tau antibodies. The neuropathological diagnosis of MSA required the presence of glial cytoplasmic inclusions.^42^ The Sydney Brain Bank provided tissue from one control patient. Demographic information about the patient samples used is shown in Supplementary Table 2.

### Extraction of α-synuclein filaments

Sarkosyl-insoluble material was extracted from fresh-frozen anterior internal capsule tissue from the MSA14 sample using a protocol similar to previous α-synuclein filament purifications.^20^ Tissue was homogenized in 20 volumes (v/w) of extraction buffer consisting of 10 mM Tris-HCl pH 7.5, 0.8 M NaCl, 10% sucrose and 1 mM EGTA. Homogenates were brought to 2% sarkosyl and incubated at 37 °C for 30 min. After centrifugation at 10,000 x *g* for 10 min, the supernatants were collected, and the pellets were resuspended in 20 volumes (v/w) of extraction buffer with 2% sarkosyl. The resuspended pellets were spun at 10,000 x *g* for 10 min once again, and the supernatants were again collected and added to the pooled supernatants from the previous step. The supernatant pool was divided and spun at 100,000 x *g* for 30 min at 22 °C. The pellets were then resuspended in 500 μL/g extraction buffer, combined, and centrifuged at 3,000 x *g* for 5 min. The supernatants were diluted three-fold in 50 mM Tris-HCl pH 7.5, 0.15 M NaCl, 10% sucrose and 0.2% sarkosyl, and spun at 166,000 x *g* for 40 min at 22 °C. The final sarkosyl-insoluble pellet was resuspended in 100 μL/g of 50 mM Tris-HCl, pH 7.5. The resuspended pellet from each preparation was centrifuged at 3,000 x *g* for 5 min to pellet any insoluble debris, and the supernatant was carried forward for negative stain and cryo-EM experiments.

To extract filaments from TgM83^+/-^ mouse tissue, fresh-frozen hindbrains from two clinical mice were combined, and the protocol above was used to extract the sarkosyl insoluble filaments. Similarly, two hindbrains collected from spontaneously ill TgM83^+/+^ mice were also combined, and the same protocol was used to extract fibrils.

### Negative stain electron microscopy

Sarkosyl-insoluble preparations were diluted 1:10 (human) and 1:5 (mouse), and 5 μL of sample was added to a glow-discharged formvar-coated 300 mesh copper grid covered with a layer of amorphous carbon. After 30 s, the grid was blotted with filter paper, washed with water, and blotted again. The wash step was repeated once more for a total of two washes before 5 μL of 0.75% uranyl formate was added and blotted after approximately 40 s. Another 5 μL aliquot of uranyl formate was added, then removed by gradual vacuum aspiration, and this was repeated once more for a total of 3 additions of uranyl formate. Images were collected on a Talos L120C (Thermo Fisher Scientific) operating at 120 kV and equipped with a Ceta-D (Thermo Fisher Scientific) camera.

### Cryo-EM grid preparation and data collection

Sarkosyl-insoluble preparations were applied to holey carbon grids (Quantifoil Au R2/2, 200 mesh) coated with a 2 nm thick layer of amorphous carbon. Mouse preparations were applied undiluted, and human preparations were diluted 5:12 before application. Aliquots of 3 μL were applied to the grids at 100% humidity and 4 °C for 15 s and blotted for 4 s before being plunge-frozen in liquid ethane using a Vitrobot Mark IV System (Thermo Fisher Scientific). Movies were collected on a 300 KeV Titan Krios G4 microscope (Thermo Fisher Scientific) equipped with a Falcon 4i direct electron detector (Thermo Fisher Scientific) in counting mode, at a nominal magnification of 165,000x resulting in a pixel size of 0.74 Å/pixel after calibration to a 1.3 Å resolution apoferritin structure solved on the same system. The total dose of each exposure averaged 45 electrons/Å^2^ and 5 electrons/pixel/s for a total of a 5.2 s exposure with a Selectris energy filter (Thermo Fisher Scientific) operating with a slit width set to 10 eV. Movies were saved in tiff format and the 1,664 raw frames were manually fractionated into 90 fraction stacks.

### Image processing

All datasets were processed in RELION using standard helical reconstruction.^43,44^ Movies were motion-corrected using MotionCor2.^45^ The dose-weighted summed micrographs were directly imported to RELION and used for further processing.^44^ Contrast transfer function (CTF) was estimated using CTFFIND 4.1.^46^ Filaments were picked manually and extracted with a box size of 1080 pixels downsampled to 360 pixels. 2D classification was used to assess data quality and filament heterogeneity. For the human MSA sample, MSA Type I_1_ (EMD-10650)^20^ and Type II_1_ (EMD-10651)^20^ maps were used as 3D initial models. For the TgM83^+/-^ sample, a 3D initial model was created de novo from the 2D class averages using RELION’s relion_helix_inimodel2d feature, using a crossover distance of 525 Å estimated from the 2D class averages. For both human and mouse samples, viable particles selected from the initial 2D class averages were re-extracted using a 288-pixel box without downsampling, and 3D classification was performed with the initial models low-pass filtered to 15 Å. The best classes were selected from the 3D classification, and the final rise and twist was optimized in 3D auto-refinement. Finally, CTF refinement was used to correct for asymmetrical and symmetrical aberrations, estimate anisotropic magnification, and refine per-particle defocus values. The final 3D maps were sharpened using standard postprocessing methods in RELION, and the final resolution was determined from Fourier shell correlation (FSC) at 0.143 Å from two independently refined half-maps using a soft-edged solvent mask. The final resolution of the human MSA filament was 2.6 Å, and the resolution of the TgM83^+/-^ filament was 2.8 Å.

### Model building and refinement

Published MSA Type I_1_ coordinates^20^ were docked into the density map of the human MSA filaments. The model was refined against the density using Isolde.^47^ For the TgM83^+/-^ filaments, coordinates of the IIB protofilament of human MSA Type II_1_ were docked into the density map. The initial model was refined against the density using Phenix^48^ and further refined in Isolde.^47^ Figures were prepared using ChimeraX.^49^

### Mouse inoculations

All mice were group housed in ABSL-2 conditions, unless health concerns deemed separation necessary. Animals were maintained on a 12 h light/dark cycle with free access to food and water in an AAALAC-accredited facility. TgM83^+/-^ mice were generated by breeding TgM83^+/+^ male mice (Strain: 004479) with B6C3F1 female mice (Strain: 100010), both of which were purchased from The Jackson Laboratory.

For TgM83^+/-^ inoculation studies, fresh-frozen C3 and MSA14 patient samples were homogenized in calcium- and magnesium-free 1X Dulbecco’s phosphate-buffered saline (DPBS; Gibco) using the Omni Tissue Homogenizer (Omni International) to create a 10% (w/v) homogenate. Total protein in each homogenate was measured via bicinchoninic acid (BCA) assay (Pierce) and adjusted to 1 mg/mL in 1X DPBS. Six-week-old TgM83^+/-^ mice were anesthetized with isoflurane prior to inoculating 30 μL into the right parietal lobe. Mice were assessed three times each week for the onset of neurological signs, including hindlimb clasping, loss of forepaw strength, ataxia, kyphosis, loss of bladder control as exhibited by urine scald, etc. Following onset of progressive neurological signs, or at experimental end point, mice were euthanized and the brain was removed. Upon removal, samples used for cryo-EM analysis were dissected to separate the hindbrain from the forebrain, and the majority of MSA-induced pathology in TgM83^+/-^ mice is in the hindbrain.^29^ All other brain samples were bisected along midline with one half fixed in formalin for neuropathological analysis and the other half frozen for cell assay and biochemical assessment.

As TgM83^+/+^ mice develop spontaneous disease,^24^ after the male breeders were retired from the colony, they were monitored three times each week for the onset clinical signs, as done with the inoculated TgM83^+/-^ mice. Following onset of progressive clinical disease, mice were euthanized and brains were collected as described above. The female B6C3F1 breeders were also aged for 1 y after breeding, at which point they were euthanized and brains were collected half-fixed and half-frozen.

### Mouse immunohistochemistry and neuropathology

The formalin-fixed mouse samples were coronally sectioned before processing through graded alcohols, xylene, and paraffin, and embedded as previously described.^29^ Eight-micron sections were mounted and deparaffinized prior to antigen retrieval (incubation in 0.1 *M*, pH 6 citrate buffer for 20 min). Slides were blocked in 10% (v/v) normal goat serum before incubating with the primary antibody EP1536Y (1:1,000; Abcam) at room temperature overnight. Primary antibodies were detected using an Alexa Fluor 488-conjugated antibody (1:1,000; A-11008; Invitrogen). Nuclei were labeled with Hoechst 33342 (1:5,000; Thermo Fisher Scientific). Slides were imaged using the Lionheart FX automated fluorescent microscope (Agilent) with the 10x objective. Images were analyzed using the BioTek Gen5 software package (Agilent; version 3.16), quantifying α-synuclein pathology in the hippocampus (HC), thalamus (Thal), hypothalamus (HTH), midbrain (Mid), and hindbrain (Hind) by setting a pixel intensity threshold based on positive and negative control slides. Representative images of mouse hindbrain pathology were acquired with the DMi8 inverted fluorescent microscope (Leica) and processed using LAS X software (Leica).

### Cellular assay for α-synuclein strain typing

Monoclonal HEK293T cells expressing chimeric human-mouse α-synuclein (human mouse with the A53T and S87N substitutions) were generated as described.^30^ These cells, along with previously developed monoclonal HEK293T cell lines expressing other mutant human α-synuclein sequences fused to YFP (α-syn-YFP)^30^ were maintained in a humid environment at 37 °C and 5% CO_2_ and cultured in Dulbecco’s modified Eagle’s medium (Corning) with 10% premium fetal bovine serum (Gibco) and 50 units/mL penicillin-streptomycin (Gibco). Cells were plated in a black walled 384-well plate (Greiner) with Hoechst 33342 (0.013 mg/well in dimethyl sulfoxide; Invitrogen) using the conditions reported in Supplementary Table S3. Frozen brain tissue samples were homogenized in 1X DPBS (10% w/v) and aggregated α-synuclein was isolated using sodium phosphotungstic acid (NaPTA; Sigma) precipitation, as previously reported.^26^ The resulting protein pellets were diluted in 1X DPBS (cell line-specific dilutions in Supplementary Table S3) before incubating with Lipofectamine 2000 (Invitrogen) at room temperature for 1.5 h. Warmed OptiMEM (Thermofisher) was added to samples prior to plating in 6 technical replicate wells per sample and the plate was subsequently covered with an adhesive membrane and incubated at 37 °C and 5% CO_2_ for 4 d. The chimeric human-mouse α-synuclein cell line was plated as described above, but in four technical replicates with a MicroPro 300 (Rainin). The Lionheart FX was used to collect DAPI and YFP images from four regions of interest per well. The BioTek Gen5 software was used to build custom algorithms to quantify the total fluorescence summed across all aggregates normalized to total cell count (x 10^5^ arbitrary units, A.U.). Values from the four regions of interest were summed to determine a single value for each well, and the replicate well values were averaged to calculate α-synuclein prion infectivity for each sample.

### Proteinase K and thermolysin digestion

Brain homogenates from TgM83^+/+^ mice were assessed for proteinase K (PK; Thermo Fisher Scientific) and thermolysin (TL; Sigma) resistance using a recently reported protocol.^16^ Briefly, 10% brain homogenates were incubated with 10x detergent extraction buffer [5% (w/v) sodium deoxycholate, 5% (v/v) NP-40] with Pierce Universal Nuclease (Thermo Fisher Scientific) and Halt Phosphatase Inhibitor Cocktail (for TL digestion only; Thermo Fisher Scientific) for 20 min on ice. Samples were centrifuged at 1,500 x *g* for 5 min, and the supernatants were collected. After measuring total protein in the supernatant by BCA assay (Pierce), samples were normalized to 500 μg total protein for PK digestion and 400 μg total protein for TL digestion. For PK digestion, samples were digested with 100 μg/mL PK in a thermomixer (Eppendorf) at 37 °C at 600 rpm for 1 h. The digestion was halted by adding 200 mM phenylmethylsulfonyl fluoride (PMSF; Sigma) prior to centrifugation at 100,000 x *g* at 4 °C for 1 h. The supernatant was removed and pellets were resuspended in 1X NuPAGE LDS sample buffer (Thermo Fisher Scientific) with 2.5% ß-mercaptoethanol (VMR Life Sciences). For TL digestion, samples were digested with 50 μg/mL TL (or same volume of buffer) in a thermomixer (Eppendorf) at 37 °C at 600 rpm for 1 h. The digestion was halted by adding 0.25 *M* EDTA (Invitrogen) prior to centrifugation at 100,000 x *g* at 4 °C for 1 h. The supernatant was removed and pellets were resuspended in the same sample buffer. For both protocols, samples were then boiled at 100 °C for 10 min.

### Immunoblotting

The PK and TL-digested samples were loaded onto a 10% Bis-Tris gel (Thermo Fisher Scientific), and SDS-PAGE were performed using the MES buffer system followed by protein transfer to a polyvinylidene fluoride membrane. The membranes were then fixed in 0.4% formaldehyde for 30 min at room temperature prior to incubating another 30 min in blocking buffer [5% (w/v) nonfat milk in 1X Tris-buffered saline with 0.05% (v/v) Tween 20] at room temperature. Membranes were probed for total α-synuclein with Clone 42 primary antibody for PK-digested samples (1:10,000; #610786; BD Biosciences) and EP1536Y for TL-digested samples (1:10,000; Abcam) in blocking buffer overnight at 4 °C in a heat-sealed pouch. Membranes were washed in 1X TBST three times prior to incubating with goat anti-mouse (PK) or goat anti-rabbit (TL) secondary antibody conjugated to horseradish peroxidase diluted in blocking buffer (1:10,000; Abcam) for 1 h. Membranes were again washed three times in 1X TBST before imaging using either the Azure 600 (Azure Biosystems) for the PK digested samples or the iBright FL1500 (Thermo Fisher Scientific) for the TL digested samples.

### Statistical analysis

Data are presented as mean ± standard deviation. Data were analyzed using GraphPad Prism software (version 10). Kaplan-Meier curves were analyzed using a log-rank Mantel-Cox test. Inoculated TgM83^+/-^ neuropathology data were analyzed using a two-way ANOVA with Šídák’s multiple comparison test. Cell assay data comparing human patient samples C3 and MSA14, as well as comparing passaged C3 with passaged MSA14, were analyzed using an unpaired *t* test with Welch’s correction. Cell assay data comparing B6C3F1 samples with TgM83^+/+^ samples were analyzed using a one-way ANOVA with Dunnett’s multiple comparison test. Significance was determined with a *P*-value < 0.05.

## ACKNOWLEDGMENTS

We thank the Colorado State University Lab Animal Resources team for their support caring for the mice used in these studies. This work was made possible by donated human patient tissue, and we would like to recognize and thank the patients and their families for this generous gift. Fig. 1a was created using BioRender.com

## AUTHOR CONTRIBUTIONS

M.M., C.R.K, G.E.M., and A.L.W. conceptualized the study. M.M., C.R.K., G.Z., G.E.M., and A.L.W. designed the experiments. M.M., A.A.M., E.T., and G.E.M. performed cryo-EM sample preparation, data collection, and data processing. M.M. and G.E.M. generated atomic models. C.R.K., M.P.F., S.A.M.H., M.D., and A.L.W. conducted animal studies and isolated sample tissues. C.R.K., G.Z., P.R., and S.A.M.H. performed neuropathological, biological, and biochemical analyses of samples. M.M., C.R.K., G.Z., G.E.M., and A.L.W. wrote the manuscript. All authors contributed to editing the manuscript.

## MATERIALS & CORRESPONDENCE

Address request for materials to Amanda L. Woerman. Send correspondence to Gregory E. Merz and Amanda L. Woerman.

## REFERENCES

1 Bessen, R. A. & Marsh, R. F. Identification of two biologically distinct strains of transmissible mink encephalopathy in hamsters. J. Gen. Virol. 73, 329–334 (1992).

2 Bessen, R. A. & Marsh, R. F. Biochemical and physical properties of the prion protein from two strains of the transmissible mink encephalopathy agent. J. Virol. 66, 2096–2101 (1992).

3 Bartz, J. C. in Prion Diseases Cold Spring Harb. Perspect. Med. (ed S.B. Prusiner) 31–44 (Cold Spring Harbor Laboratory Press, 2017).

4 Watts, J. C. et al. Transmission of multiple system atrophy prions to transgenic mice. Proc Natl Acad Sci U S A 110, 19555–19560 (2013).

5 Prusiner, S. B. et al. Evidence for α-synuclein prions causing multiple system atrophy in humans with parkinsonism. Proc. Natl. Acad. Sci. U.S.A. 112, E5308–E5317 (2015).

6 Woerman, A. L. et al. Familial Parkinson’s point mutation abolishes multiple system atrophy prion replication. Proc. Natl. Acad. Sci. USA 115, 409–414 (2018).

7 Woerman, A. L. et al. Multiple system atrophy prions retain strain specificity after serial propagation in two different Tg(*SNCA*\*A53T) mouse lines. Acta Neuropathol. 137, 437–454 (2019).

8 Ayers, J. I. et al. Different α-synuclein prion strains cause dementia with Lewy bodies and multiple system atrophy. Proc Natl Acad Sci U S A 119, e211349119 (2022).

9 Holec, S. A. M. et al. The E46K mutation modulates α-synuclein prion replication in transgenic mice. PLoS Pathog 18, e1010956 (2022).

10 Holec, S. A. M. et al. Multiple system atrophy prions transmit neurological disease to mice expressing wild-type human α-synuclein. Acta Neuropathol 144, 677–690 (2022).

11 Holec, S. A. M., Khedmatgozar, C. R., Schure, S. J., Pham, T. & Woerman, A. L. A-synuclein prion strains differentially adapt after passage in mice. PLoS Pathog 20, e1012746 (2024).

12 Holec, S. A. M. et al. Evidence of a novel alpha-synuclein strain isolated from a Parkinson’s disease with dementia patient sample. Acta Neuropathol Commun 13 (2025).

13 Holec, S. A. M., Khedmatgozar, C. R., Schure, S. J., Bartz, J. C. & Woerman, A. L. Co-infection with two alpha-synuclein strains reveals novel synergistic interactions. Acta Neuropathol 150, 50 (2025).

14 Yamasaki, T. R. et al. Parkinson’s disease and multiple system atrophy have distinct α-synuclein seed characteristics. J. Biol. Chem. 294, 1045–1058 (2019).

15 Lau, A. et al. alpha-Synuclein strains target distinct brain regions and cell types. Nat Neurosci 23, 21–31 (2020).

16 So, R. W. L. et al. Stochastic misfolding drives the emergence of distinct α-synuclein strains. Neuron S0896-6273, 00040–00041 (2026).

17 Shahnawaz, M. et al. Discriminating alpha-synuclein strains in Parkinson’s disease and multiple system atrophy. Nature 578, 273–277 (2020).

18 Strohaker, T. et al. Structural heterogeneity of alpha-synuclein fibrils amplified from patient brain extracts. Nat Commun 10, 5535 (2019).

19 Ayers, J. I. et al. Localized Induction of Wild-Type and Mutant Alpha-Synuclein Aggregation Reveals Propagation along Neuroanatomical Tracts. J Virol 92 (2018).

20 Schweighauser, M. et al. Structures of α-synuclein filaments from multiple system atrophy. Nature 585, 464–469 (2020).

21 Yan, N. L. et al. Cryo-EM structure of a novel alpha-synuclein filament subtype from multiple system atrophy. FEBS Lett 599, 33–40 (2025).

22 Yang, Y. et al. Structures of alpha-synuclein filaments from human brains with Lewy pathology. Nature 610, 791–795 (2022).

23 Yang, Y. et al. New SNCA mutation and structures of α-synuclein filaments from juvenile-onset synucleinopathy. Acta Neuropathol 145, 561–572 (2023).

24 Giasson, B. I. et al. Neuronal α-synucleinopathy with severe movement disorder in mice expressing A53T human α-synuclein. Neuron 34, 521–533 (2002).

25 Holec, S. A. M. & Woerman, A. L. Evidence of distinct α-synuclein strains underlying disease heterogeneity. Acta Neuropathol 142, 73–86 (2021).

26 Woerman, A. L. et al. Propagation of prions causing synucleinopathies in cultured cells. Proc. Natl. Acad. Sci. U.S.A. 112, E4949–E4958 (2015).

27 Burger, D. et al. Synthetic α-synuclein fibrils replicate in mice causing MSA-like pathology. Nature 648, 409–417 (2025).

28 Zhang, H. et al. Dimerisation and twist reversal of the Lewy fold in a-synuclein mutants with Parkinson’s disease and dementia. bioRxiv 04.07.716933 (2026).

29 Woerman, A. L. et al. Kinetics of α-synuclein prions preceding neuropathological inclusions in multiple system atrophy. PLOS Pathogens 16, e1008222 (2020).

30 Reis, P. M. et al. Structurally targeted mutagenesis identifies key residues supporting α-synuclein misfolding in multiple system atrophy. J Parkinsons Dis 14, 1543–1558 (2024).

31 Taylor, R. Short Nonbonded Contact Distances in Organic Molecules and Their Use as Atom-Clash Criteria in Conformer Validation and Searching. J Chem Inf Model 51, 897–908 (2011).

32 Bernis, M. E. et al. Prion-like propagation of human brain-derived alpha-synuclein in transgenic mice expressing human wild-type alpha-synuclein. Acta Neuropathol. Commun. 3, 75 (2015).

33 Woerman, A. L. et al. MSA prions exhibit remarkable stability and resistance to inactivation. Acta Neuropathol. 135, 49–63 (2018).

34 Breid, S. et al. Neuroinvasion of α-synuclein prionoids after intraperitoneal and intraglossal inoculation. J. Virol. 90, 9182–9193 (2016).

35 Wang, F. et al. Seed amplification of MSA alpha-synuclein aggregates preserves the biological and structural properties of brain-derived aggregates. Nat Commun 16, 11266 (2025).

36 Yan, N. L. et al. Cryo-EM structure of a novel α-synuclein filament subtype from multiple system atrophy. FEBS Lett 599, 33–40 (2025).

37 Luk, K. C. et al. Molecular and Biological Compatibility with Host Alpha-Synuclein Influences Fibril Pathogenicity. Cell Rep 16, 3373–3387 (2016).

38 Kraus, A. et al. High-resolution structure and strain comparison of infectious mammalian prions. Mol Cell 81, 4540–4551 (2021).

39 Prusiner, S. et al. Transgenetic studies implicate interactions between homologous PrP isoforms in scrapie prion replication. Cell 63, 673–686 (1990).

40 Ghaemmaghami, S., May, B. C. H., Renslo, A. R. & Prusiner, S. B. Discovery of 2-aminothiazoles as potent antiprion compounds. J. Virol. 84, 3408–3412 (2010).

41 Berry, D. B. et al. Drug resistance confounding prion therapeutics. Proc. Natl. Acad. Sci. U.S.A. 110, E4160–E4169 (2013).

42 Gilman, S. et al. Second consensus statement on the diagnosis of multiple system atrophy. Neurology 71, 670–676 (2008).

43 He, S. & Scheres, S. H. W. Helical reconstruction in RELION. J Struct Biol 198, 163–176 (2017).

44 Kimanius, D., L., D., Sharov, G., Nakane, T. & Scheres, S. H. W. New tools for automated cryo-EM single-particle analysis in RELION-4.0. Biochem J 478, 4169–4185 (2021).

45 Zheng, S. Q. et al. MotionCor2: anisotropic correction of beam-induced motion for improved cryo-electron microscopy. Nat Methods 14, 331–332 (2017).

46 Rohou, A. & Grigorieff, N. CTFFIND4: Fast and accurate defocus estimation from electron micrographs. J Struct Biol 192, 216–221 (2015).

47 Croll, T. I. ISOLDE: a physically realistic environment for model building into low-resolution electron-density maps. Acta Crystallogr D Biol Crystallogr 74, 519–530 (2018).

48 Adams, P. D. et al. PHENIX: a comprehensive Python-based system for macromolecular structure solution. Acta Crystallogr D Biol Crystallogr 66, 213–221 (2010).

49 Pettersen, E. F. et al. UCSF ChimeraX: Structure visualization for researchers, educators, and developers. Protein Sci 30, 70–82 (2021).

