## Supplemental Material for "Adaptation of α-synuclein fibrils following multiple system atrophy transmission to mice" (from another version of this preprint: version 1, 2026-05-11)

[‡]Current affiliation: Department of Neuroscience, UConn Health, Farmington, CT, USA.

[¶]Current affiliation: SiVEC Biotechnologies, Fort Collins, CO, USA.

### SUPPLEMENTARY FIGURES

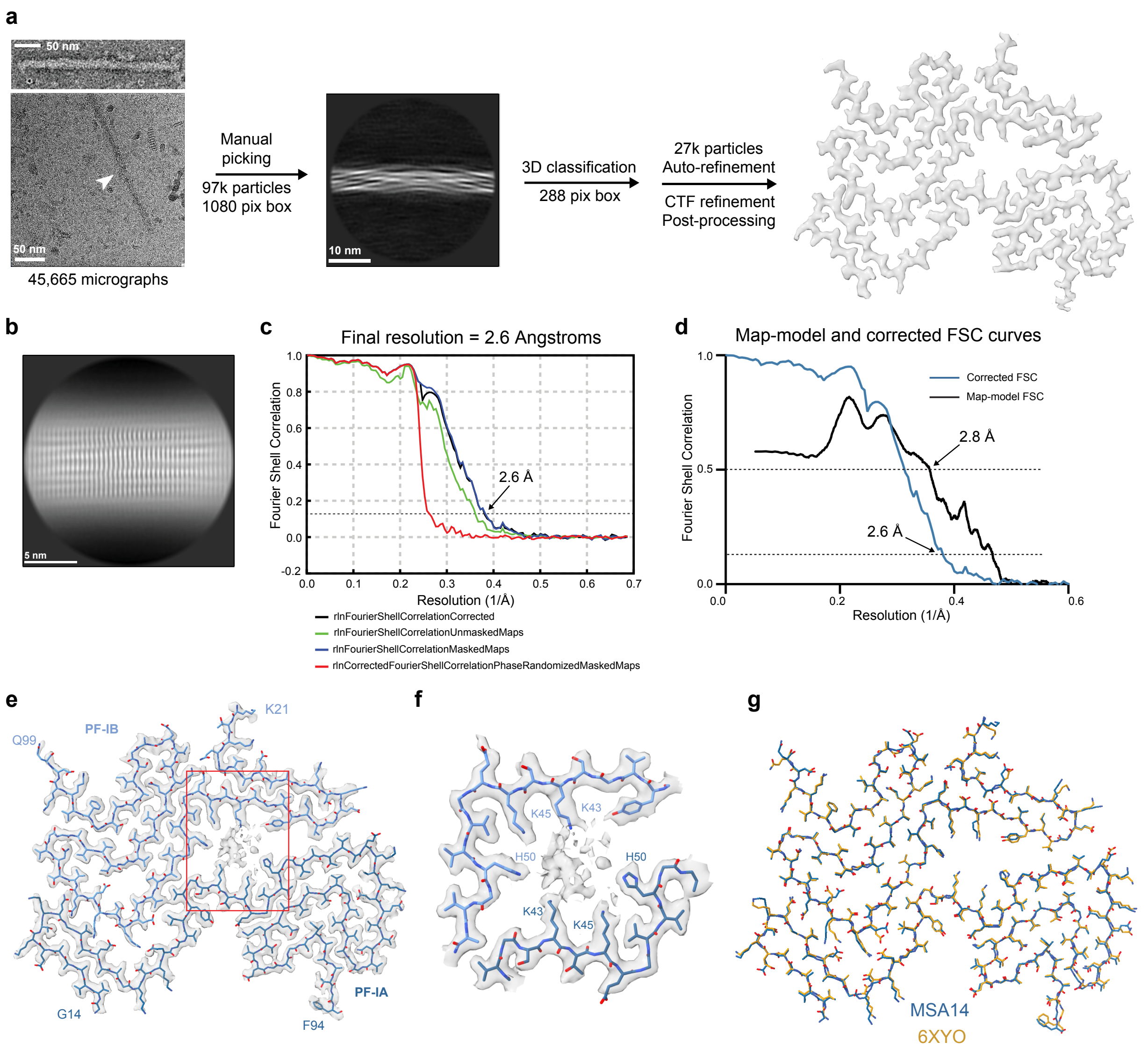

**Supplementary Figure 1. Resolution of α-synuclein fibrils from patient sample MSA14. a**, Processing workflow for α-synuclein filaments purified from MSA14. Representative negative stain TEM (top left) and cryo-EM micrograph (bottom left) with representative 2D class average (middle, 785 Å box downsampled three-fold). Final cryo-EM map on the right, resolved to 2.6 Å. **b**, 2D class average of unbinned particles, with a box size of 210 Å. **c**, Fourier shell correlation (FSC) curves of two independently refined half-maps of the MSA14 fold. **d**, Corrected final map (blue) and map-model (black) FSC curves. **e**, Cryo-EM structure of α-synuclein filaments purified from the anterior internal capsule of patient MSA14. The atomic model (blue) fit into the cryo-EM density map (grey) with non-proteinaceous density identified by the red box. **f**, Inset of the non-proteinaceous density at the protofilament interface of the MSA14 cryo-EM structure. **g**, Alignment of our MSA14 model with the previously published model (PDB: 6XYO[1], RMSD = 0.503 Å).

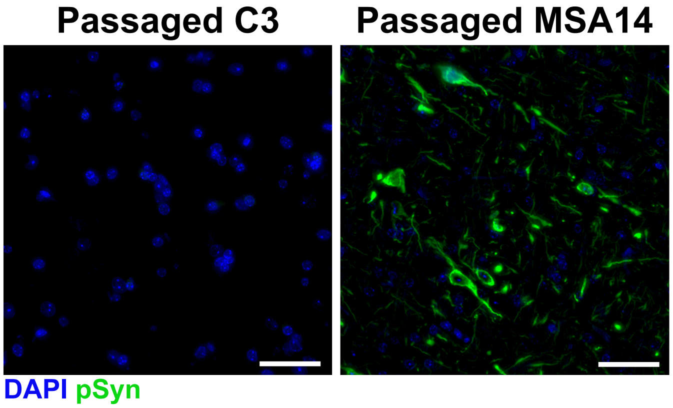

**Supplementary Figure 2. Representative neuropathology in TgM83$^{+/-}$ mice.** Representative images of phosphorylated α-synuclein neuropathology (green; EP1536Y; 1:1,000) in the hindbrain of TgM83$^{+/-}$ mice inoculated with either C3 (left) or MSA14 (right) patient sample. DAPI shown in blue. Scale bar, 50 μm.

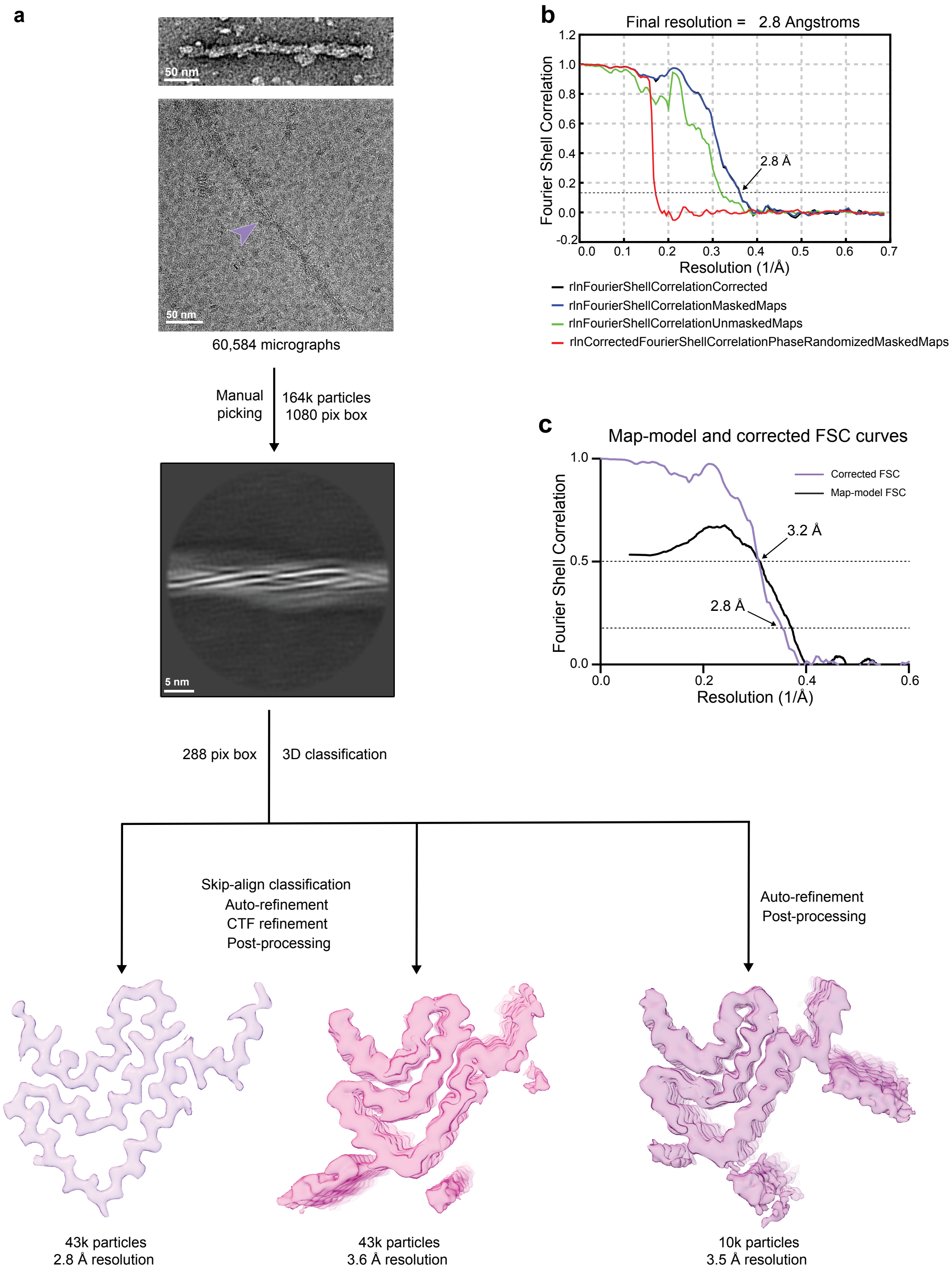

**Supplementary Figure 3. Resolution of α-synuclein fibrils from TgM83$^{+/-}$ mice inoculated with patient sample MSA14. a**, Processing workflow of mouse-passaged MSA filaments. Representative negative stain TEM and cryo-EM micrograph (top) with representative 2D class average (middle, 785 Å box downsampled three-fold) of sarkosyl-insoluble α-synuclein filaments. Highest-resolution cryo-EM map (bottom left), with classification yielding lower-resolution maps with additional densities extending from the TgM83$^{+/-}$-passaged MSA14 α-synuclein fibrils (middle and right, bottom). **b**, Fourier shell correlation (FSC) curves of two independently refined half-maps of the MSA14 fold. **c**, Corrected final map (purple) and map-model (black) FSC curves.

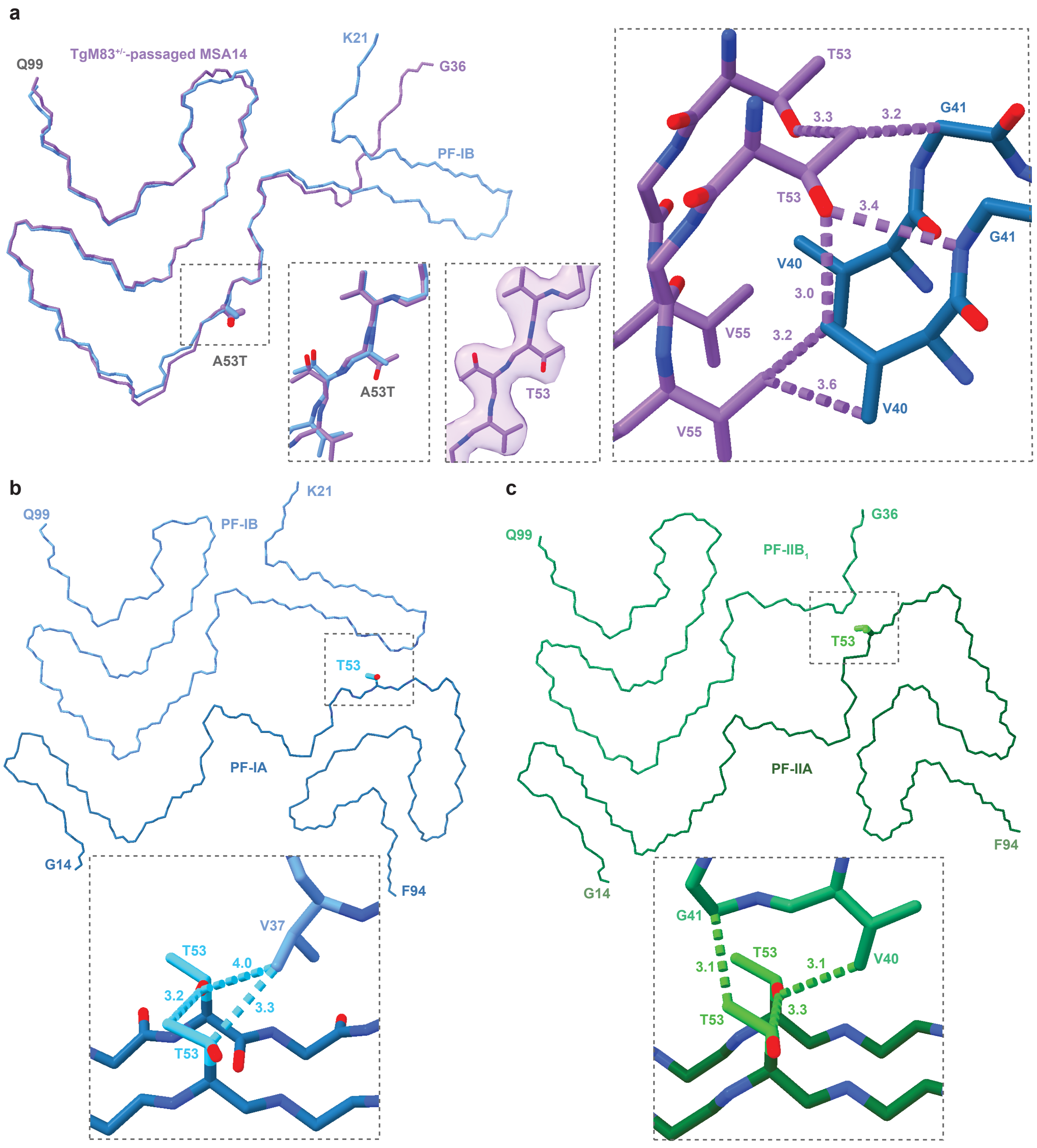

**Supplementary Figure 4. The A53T mutation does not alter MSA14 replication in TgM83$^{+/-}$ mice. a**, Alignment of the mouse-passaged MSA14 structure (purple) with PF-IB of the MSA14 fold (blue), showing T53 is unperturbed relative to A53 in the human fold (left inset). The well-defined density for the mouse-passaged T53 side chain is shown in the middle inset. In the right inset, the side chains of T53 of the mouse-passaged fold and V40 of the aligned PF-IA of MSA14 adopt conformations that satisfy steric requirements. **b & c**, Modeling the A53T mutation in (**b**) MSA PF-IA (PDB: 6XYO) and (**c**) MSA PF-IIA (PDB: 6XYP) fibrils reported by Schweighauser *et al.*[1] shows a threonine substitution is tolerated in these structures.

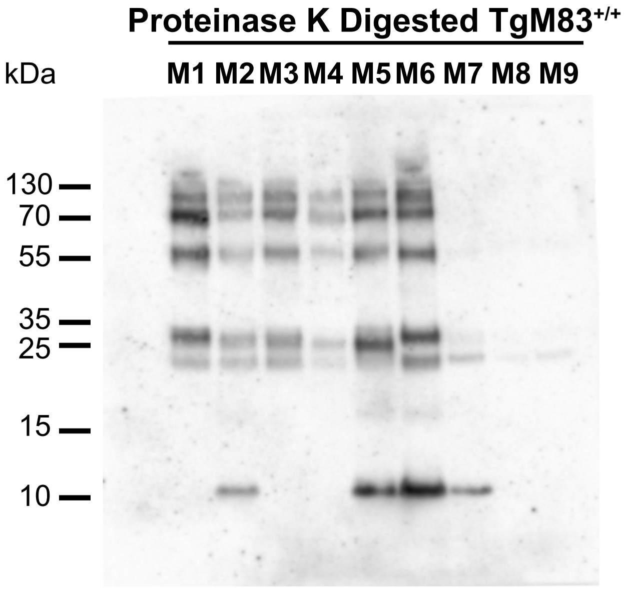

**Supplementary Figure 5. Uncropped Proteinase K digestion blot.** Uncropped copy of the Western blot in Fig. 5c probing for total insoluble α-synuclein (Clone 42; 1:10,000) following proteinase K digestion of brain homogenates from spontaneously ill TgM83$^{+/+}$ mice.

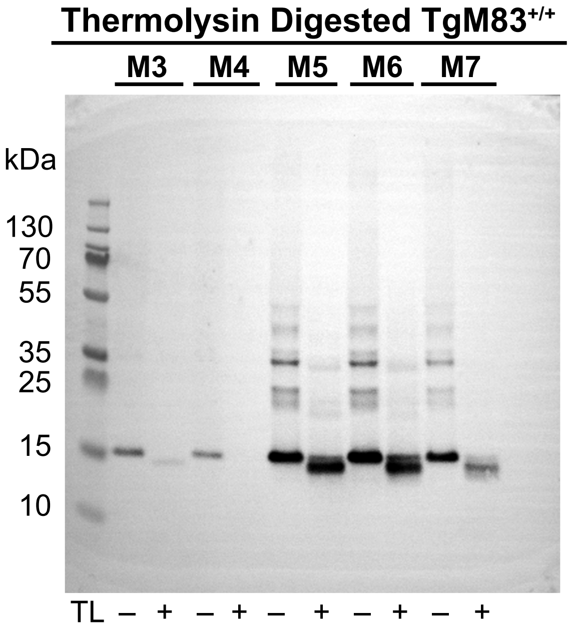

**Supplementary Figure 6. Uncropped thermolysin digestion blot.** Uncropped copy of the Western blot in Fig. 5d probing for total insoluble α-synuclein (Clone 42; 1:10,000) with and without thermolysin (TL) digestion of brain homogenates from spontaneously ill TgM83$^{+/+}$ mice.

### SUPPLEMENTAL TABLES

**Supplementary Table 1. Cryo-EM data collection and model refinement statistics.**

| | Human MSA14 (EMDB-XXXXX) (PDB YYYY) | TgM83$^{+/-}$-passaged MSA14 (EMDB-XXXX) (PDB YYYY) |
|---|---|---|
| **Data collection and processing** | | |
| Magnification | 165,000 | 165,000 |
| Voltage (kV) | 300 | 300 |
| Electron exposure (e–/Å$^2$) | 45 | 45 |
| Defocus range (μm) | -0.8 to -1.2 | -0.8 to -1.2 |
| Pixel size (Å) | 0.728 | 0.278 |
| Symmetry imposed | C1 | C1 |
| Initial particle images (no.) | 97,491 | 164,763 |
| Final particle images (no.) | 27,285 | 43,393 |
| Map resolution (Å) | 2.6 | 2.8 |
| FSC threshold | 0.143 | 0.143 |
| Map resolution range (Å) | 10-2.2 | 10-2.5 |
| **Refinement** | | |
| Initial model used (PDB code) | 6XYO | De novo |
| Model resolution (Å) | 2.8 | 3.2 |
| FSC threshold | 0.5 | 0.5 |
| Model resolution range (Å) | 19.1-2.1 | 17.5-2.5 |
| Map sharpening *B* factor (Å$^2$) | -58.8 | -64.7 |
| Model composition | | |
| Non-hydrogen atoms | 5,490 | 2,220 |
| Protein residues | 800 | 320 |
| Ligands | 0 | 0 |
| *B* factors (Å$^2$) | | |
| Protein | 45.09 | 113.99 |
| Ligand | N/A | N/A |
| R.m.s. deviations | | |
| Bond lengths (Å) | 0.010 | 0.010 |
| Bond angles (°) | 1.845 | 1.852 |
| Validation | | |
| MolProbity score | 2.56 | 2.45 |
| Clashscore | 26.77 | 18.72 |
| Poor rotamers (%) | 1.85 | 2.17 |
| Ramachandran plot | | |
| Favored (%) | 83.08 | 93.55 |
| Allowed (%) | 6.92 | 4.84 |
| Disallowed (%) | 0.00 | 1.61 |

**Supplementary Table 2. Patient sample information.**

| Patient | Neurological Disease | Age at Death | Sex | Brain Region | Brain Bank |
|---|---|---|---|---|---|
| C3 | None | 84 | M | Pons | Sydney Brain Bank |
| MSA14 | MSA | 68 | M | Midbrain | MADRC[a] |

[a]Massachusetts Alzheimer's Disease Research Center

**Supplementary Table S3. A-synuclein biosensor cell assay conditions.**

| Mutation(s) | Cells Per Well | Lipofectamine (%) | Sample Dilution in DPBS |
|---|---|---|---|
| A30G | 3,000 | 1% | 1:5 |
| E46K | 3,500 | 1.5% | 1:10 |
| G51D | 3,500 | 2% | 1:4 |
| A53E | 3,500 | 1.5% | 1:10 |
| A53T | 3,500 | 1.5% | 1:14 |
| V55Y | 3,500 | 1% | 1:20 |
| V66F | 4,250 | 1% | 1:10 |
| V74P | 4,500 | 1.5% | 1:10 |
| A53T,S87N | 4,500 | 1% | 1:10 |

### SUPPLEMENTARY REFERENCE

1 Schweighauser, M. *et al.* Structures of α-synuclein filaments from multiple system atrophy. *Nature* **585**, 464–469 (2020).
